# Pretraining Objective Shapes Cross-Category Generalization in Affective Image Prediction: A Geometric Comparison of Vision Transformer Encoders

**DOI:** 10.64898/2026.05.11.724194

**Authors:** Shohei Tsuchimoto, Yuka O Okazaki, Kenichi Yuasa, Sakura Nishijima, Mebuki Izumiya, Makoto Hagihara, Ryo Fujihira, Keiichi Kitajo

## Abstract

The geometry of representations learned by deep neural networks is shaped jointly by architecture and pretraining objective, yet disentangling these two factors remains difficult. Here we isolate the contribution of pretraining objective by comparing two Vision Transformers from the same backbone family but trained under different objectives: language–image contrastive learning (CLIP) and ImageNet-21k classification. Using continuous Valence–Arousal prediction on the OASIS dataset as a probe of representational quality, we evaluated frozen features under Leave-One-Theme-Out and Leave-One-Category-Out cross-validation, the latter requiring extrapolation to entirely unseen semantic categories. The contrastively pretrained encoder generalized substantially better than the classification-pretrained encoder under both protocols, with the gap widening sharply when held-out categories required cross-category generalization. To characterize why the two representations differ, we developed a geometric analysis of prediction errors, treating per-image errors as vectors in the affective plane and quantifying their spatial structure via weighted phase-locking, trajectory-based occupancy entropy, and effective dimensionality. The classification-pretrained representation collapsed errors into a small number of attractor regions with a strong center-ward pull, whereas the language-aligned representation distributed errors broadly across the affective space. Layer-wise linear probing further revealed that affective information was distributed across depth in the contrastive encoder but increasingly concentrated in deeper layers of the classification encoder, mirroring the texture-bias and category-anchored statistics characteristic of ImageNet-trained representations. These results provide a representation-geometric account of how the choice of pretraining objective, holding architecture constant, determines whether learned features generalize across semantic boundaries or remain confined to category-bound visual regularities.

**Highlights:**

- Isolate the effect of pretraining objective by holding the Vision Transformer backbone constant.
- Contrastively pretrained features generalize across unseen semantic categories where classification-pretrained features fail.
- Introduce a geometric analysis of prediction errors based on phase-locking and occupancy entropy.
- Classification pretraining produces concentrated error attractors and a rigid centerward bias.
- Affective information is distributed across depth in CLIP but localized in late layers of the classification ViT.

## Introduction

Predicting affective responses to images, defined by the perceived pleasantness (valence) and activation (arousal) of a viewer, is essential for applications spanning content recommendation, psychological stimulus selection, and human–computer interaction (Russell, 1980; Zhao et al., 2022). Over the past decade, deep vision encoders have increasingly been adopted as backbones for feature extraction, often paired with a lightweight regression head while keeping the encoder weights frozen to preserve pretrained representations (e.g., Mertens et al., 2024). While much of this progress has been driven by architectural improvements, from convolutional neural networks (CNNs) to Vision Transformers, the role of the pretraining objective remains comparatively under-examined.

This importance stems from the fact that affective responses are primarily driven by semantic content, encompassing both the depicted objects and their subjective significance, rather than by low-level visual properties such as texture or color (Borth et al., 2013; Machajdik & Hanbury, 2010). A ViT pretrained on ImageNet for object classification (Dosovitskiy et al., 2021; Steiner et al., 2021) is optimized to discriminate visual categories, and its representations reflect that: they are organized around low-level visual features, including a known bias toward texture statistics (Geirhos et al., 2019), and encode little semantic information beyond the distinctions required for category classification. In contrast, CLIP (Radford et al., 2021), trained via contrastive learning on hundreds of millions of image–text pairs, learns a representation aligned with natural-language descriptions, including the affectively loaded vocabulary humans use to describe scenes. Whether this difference in representational structure translates into better affect prediction has not been tested under conditions that hold architecture constant. Without such control, observed performance differences cannot be unambiguously attributed to the pretraining objective rather than to architectural confounders.

Prior comparisons of encoders for affective tasks have not isolated the effect of pretraining objective from that of architecture. For example, comparing CLIP (ViT-B/32, language–image contrastive) with DINOv2 (ViT-L/14, self-supervised, Mertens et al., 2024) conflates pretraining objective with model size and patch resolution; comparisons against ResNet-based models (Bustos et al., 2023) conflate the shift from convolutional to transformer architectures with the shift from classification to contrastive pretraining. When multiple variables change simultaneously, it is unclear whether observed performance differences should be attributed to how the model was trained or to how it processes images. A further limitation of much prior work is the evaluation design: random-split cross-validation does not assess whether a model generalizes to genuinely unseen content. The Open Affective Standardized Image Set (OASIS; Kurdi et al., 2017) provides an unusual opportunity here, offering psychometrically validated, publicly available normative Valence–Arousal ratings for 900 images across four semantic categories (Object, Person, Animal, Scene), yet it remains underexplored computationally. This four-category structure is well suited to a Leave-One-Category-Out evaluation protocol, making OASIS particularly appropriate for testing whether an encoder generalizes across semantically distinct content classes rather than merely interpolating within a training distribution. Prior work predicting valence and arousal from OASIS images has relied on low-level image properties or shallow CNNs, with Valence *R*^2^ rarely exceeding 0.10 (Priyadarshani & Miyapuram, 2025; Redies et al., 2020); no study has applied transformer-based encoders or evaluated generalization by holding out entire semantic categories at test time. Beyond aggregate performance metrics, the spatial structure of prediction errors across the Valence–Arousal plane has received little attention: existing studies have not characterized where in affective space two encoders diverge, nor have they examined whether errors are concentrated in systematic directional biases or distributed broadly across the rating space.

To isolate the contribution of pretraining objective to representational quality, we utilized both encoders as frozen feature extractors, so that any observed performance difference is attributable to what was learned during pretraining rather than to task-specific adaptation. Evaluating these frozen representations under Leave-One-Theme-Out (LOTO) and Leave-One-Category-Out (LOCO) cross-validation tests whether the learned representations support generalization to thematically and categorically novel content. This property is determined at pretraining time and cannot be assessed by random-split evaluation. We addressed these gaps by comparing two Vision Transformer encoders that share the same backbone family but differ in pretraining objective: CLIP (ViT-B/32, language–image contrastive learning) and an ImageNet-21k–pretrained ViT (ViT-B/16, image classification). We addressed these gaps by comparing two Vision Transformer encoders that share the same backbone family but differ in pretraining objective: CLIP (ViT-B/32, language–image contrastive learning) and an ImageNet-21k– pretrained ViT (ViT-B/16, image classification), ensuring that performance differences reflect the pretraining objective rather than architectural capacity. The overall experimental workflow and our geometric analysis framework are summarized in Figure 1. Generalization is assessed under nested LOTO (248 folds) and LOCO (4 folds) cross-validation. To elucidate the mechanisms underlying performance differences, we apply a spatially resolved, cluster-corrected error analysis to identify regions of the Valence– Arousal plane where prediction accuracy diverges between the two encoders. Beyond these scalar assessments, we introduce a geometric framework utilizing error vector fields to characterize systematic prediction biases. Unlike conventional metrics such as MSE or *R*^2^, this approach captures the directional displacement of errors, revealing toward which affective regions the predictions are systematically pulled. This geometric characterization is further extended through trajectory-based occupancy-count simulations, which visualize the spatial structure and long-term stability of prediction biases. Finally, we employ layer-wise linear probing to determine the network depth at which affective information is most robustly represented. Together, these analyses test whether the choice of pretraining objective determines representational quality for affective image prediction under conditions that require generalization to novel semantic content.

**Figure 1.**
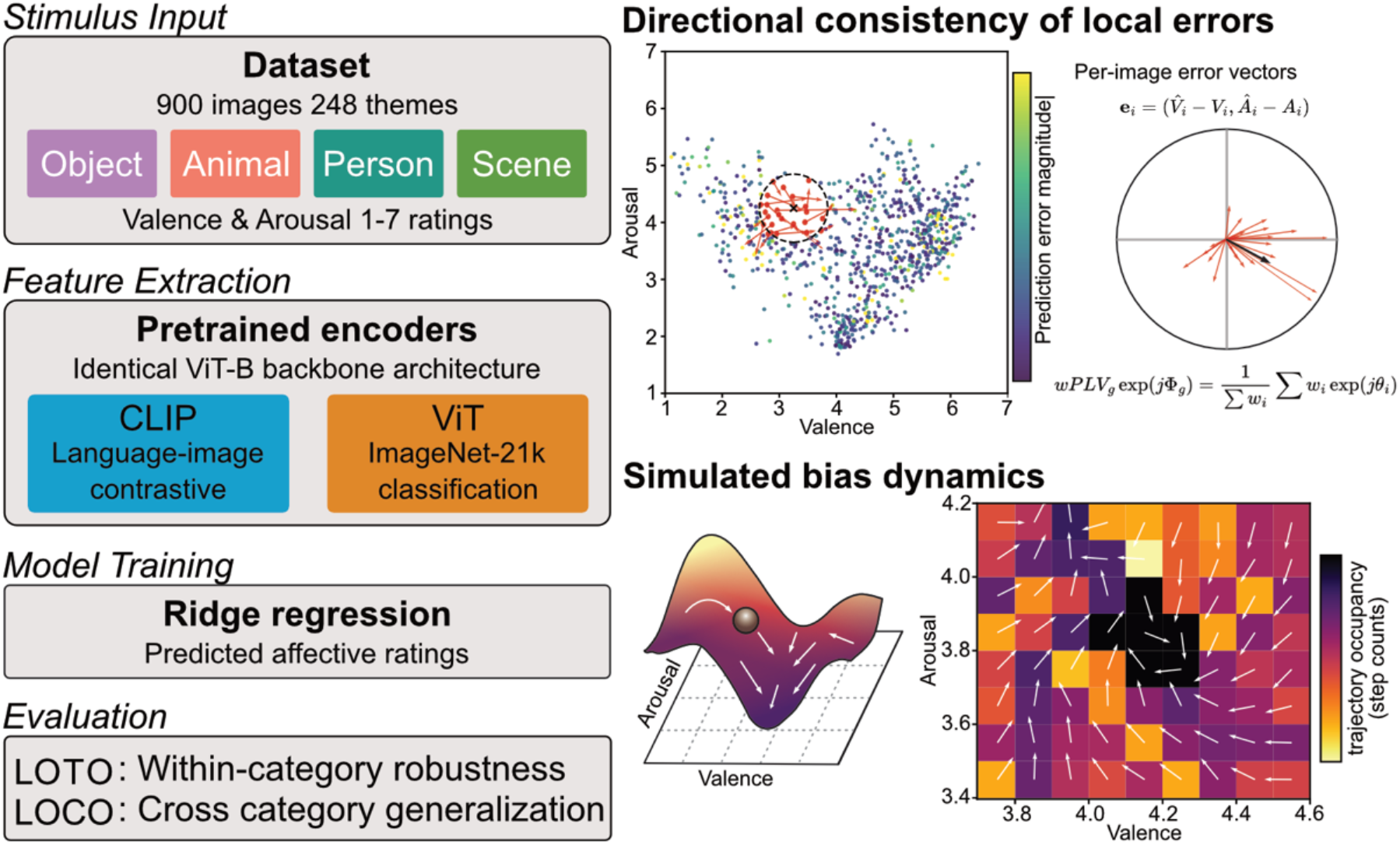
Experimental workflow and geometric analysis of systematic prediction biases. (Left) Experimental overview. OASIS images (*N*=900) were processed by two frozen Vision Transformer (ViT-B) encoders (CLIP vs. ImageNet-ViT). Valence– Arousal prediction was evaluated using Ridge regression under Leave-One-Theme-Out (LOTO) and Leave-One-Category-Out (LOCO) protocols. **(Right Top) Directional consistency of local errors**. For each image, an error vector 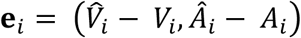, was defined, where 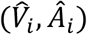 are the predicted ratings and (*V*_*i*_, *A*_*i*_) are the ground-truth ratings. These errors are visualized as a 2D scatter plot in the Valence–Arousal space, color-coded by their magnitude ( *w*_*i*_ = ‖**e**_*i*_‖ ). Local systematic bias was quantified using the weighted phase locking value (wPLV). In this calculation, **e**_*i*_ vectors were converted to phase angles (*θ*_*i*_) and magnitudes (*w*_*i*_). The representative bias direction Φ_*g*_ is determined disproportionately by larger error magnitudes *w*_*i*_, ensuring the field reflects significant predictive failures. The circular plot illustrates individual error vectors (red) and the resulting representative bias direction (black arrow). **(Right Bottom) Simulated bias dynamics**. A dynamic simulation (ball simulation) was performed by integrating the representative bias field Φ_*g*_ (white arrows). Starting from random points in the Valence–Arousal space, trajectories moved according to the local bias field, and the cumulative visit count was recorded. Regions with high visit counts, quantified in the 2D heatmap (trajectory occupancy), were projected as a simulated 3D landscape. This visualization does not reflect a static error landscape; instead, it represents emergent bias sinks (or attractors), which are regions within specific affective subspaces where predictive failures consistently concentrate.

## Methods

### Dataset

We used the OASIS datasets (Kurdi et al., 2017), a publicly available collection of 900 images with normative Valence (1–7, unpleasant to pleasant) and Arousal (1–7, calm to excited) ratings collected from a demographically diverse online sample under controlled instructions. Each image belongs to one of four semantic categories (Object, Person, Animal, Scene) and to a finer-grained semantic theme (e.g., Acorns, Alcohol, Baby; 248 themes in total), which is reflected in the individual filenames (e.g., “Acorns1.jpg”, “Acorns2.jpg”). We used the per-image mean Valence and Arousal across raters as regression targets throughout. This study used only publicly available, anonymized data and did not involve new human participants; no ethical approval was required.

### Image Encoders and Feature Extraction

We compared two frozen Vision Transformer encoders, both of which followed the Vision Transformer Base (ViT-B) architecture consisting of 12 Transformer layers. These encoders were used without any task-specific fine-tuning on the OASIS dataset. The first was the image encoder of CLIP (ViT-B/32; Radford et al., 2021), pretrained under language–image contrastive learning on 400 million image–text pairs and producing a 512-dimensional embedding per image. The second was a ViT pretrained on ImageNet-21k for image classification (ViT-B/16; Dosovitskiy et al., 2021; Hugging Face: google/vit-base-patch16-224; Wolf et al., 2020), from which we extracted the [CLS] token representation (768 dimensions). Both encoders received images resized to 224×224 and normalized with the standard ImageNet mean and standard deviation. Encoder weights were fixed; no gradient updates were applied.

We selected ViT-B/16 rather than ViT-B/32 as the classification baseline after a preliminary comparison under both single-output and joint Valence–Arousal prediction showed ViT-B/16 to yield consistently higher *R*^2^; details of that comparison are given in the Supplementary Information. The main comparison is therefore CLIP (ViT-B/32) versus ViT (ViT-B/16), with the classification-pretrained encoder given the advantage of the stronger variant; any residual performance difference can thus be attributed to pretraining objective rather than to one encoder being architecturally weaker.

### Prediction Protocol and Cross-Validation

For each encoder, we fitted a Ridge regression head to the extracted features to predict Valence and Arousal jointly as a two-output model (joint 2D prediction). In a preliminary comparison, this 2D formulation yielded higher mean *R*^2^ than separate 1D models (one Ridge per dimension); we therefore adopted 2D prediction for all main analyses. The formal comparison of 1D vs 2D prediction is detailed in the Supplementary Material (Fig. S1, Table S1). Features were standardized to zero mean and unit variance using statistics computed on the training set of each fold only; the same scaler was applied to validation and test data so that no test information was used. Targets were normalized to the range 0 to 1 by min-max scaling with fixed scale bounds (min = 1, max = 7 for both Valence and Arousal):

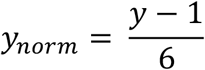

Because these bounds are fixed for the task, the same formula was applied to training, validation, and test sets without using any data-dependent statistics; predictions were back-transformed to the 1 to 7 scale only for reporting *R*^2^. We reported the coefficient of determination *R*^2^ averaged over Valence and Arousal as the primary metric. Unless otherwise noted, *R*^2^ was computed based on the aggregated test predictions across all folds (i.e., all held-out images are concatenated before computing *R*^2^ ), rather than averaging fold-level *R*^2^ values.

The regularization parameter *α* was selected under a nested procedure to ensure it never depended on test data. Within each outer training set we held out a validation split by theme (80/20) and searched *α* ∈ [0.1, 1000] in log space to maximize mean *R*^2^ on the inner validation set, using Optuna with the TPE (Tree-structured Parzen Estimator) sampler (30 trials for LOTO, 50 for LOCO). The selected *α* was then used to refit Ridge on the full outer training set before evaluating on the held-out test fold. We used two outer cross-validation protocols. LOTO held out each of the 248 semantic themes in turn, training on the remaining 247 themes; this assesses generalization to novel thematic content. LOCO held out each of the four semantic categories in turn, training on the other three; this assesses generalization across high-level content classes. For both LOTO and LOCO, permutation tests used the theme as the unit of exchange (248 themes). For LOTO we reported mean *R*^2^ with 95% bootstrap confidence intervals (5000 bootstrap samples) and Cohen’s *d* for the between-encoder difference.

### Spatially Resolved Error Analysis in Valence–Arousal Space

To examine where in affective space the two encoders differed in prediction error, we used per-image MSE values from the LOTO predictions. LOTO was used rather than LOCO because it produces a test prediction for every image, providing sufficient coverage of the Valence–Arousal space; LOCO assigns each image to only one test fold and does not yield predictions for the full image set simultaneously. We placed a regular grid on the Valence–Arousal map (step size 0.1) and at each grid point identified images whose true ratings fell within a radius of 0.5 in 1 to 7 scale. Grid points with fewer than five such images were excluded. Within each local neighborhood we performed a paired Wilcoxon signed-rank test comparing MSE_CLIP_ and MSE_ViT_; with the local effect defined as the mean of (MSE_ViT_ - MSE_CLIP_); positive values indicated lower CLIP error.

Multiple comparisons across the grid were controlled using a permutation cluster test (Maris & Oostenveld, 2007). Under each of 5000 permutations, the sign of (MSE_ViT_ - MSE_CLIP_) was flipped independently per image, local statistics were recomputed, and contiguous supra-threshold cells (uncorrected *p* < 0.05, 4-connectivity) were grouped into clusters. The null distribution of maximum cluster size was used to assign a corrected *p*-value to each observed cluster (*cluster_alpha* = 0.05). The analysis was applied over all images and separately within each category (Fig. S2).

### Error Vector Field and Dynamics Analysis

To characterize systematic prediction biases, we constructed an error vector field. For each image *i*, an error vector was defined as 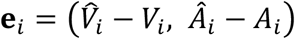, where (*V*_*i*_, *A*_*i*_) and 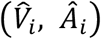, denote the ground-truth and predicted ratings for Valence and Arousal, respectively. At each grid point *g*, a local bias was calculated from neighboring images within a radius of 0.5 in 1 to 7 scale. To emphasize regions with high directional consistency, we employed the weighted phase locking value (wPLV), defined as:

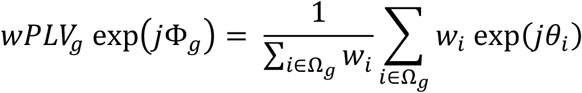

where *w*_*i*_ = ‖**e**_*i*_‖ represents the magnitude of the error vector, *θ*_*i*_ is its phase angle, and Ω_*g*_ denotes the set of images within the local neighborhood. The resulting phase angle Φ_*g*_ was utilized as the representative bias direction at each grid point, while the *wPLV*_*g*_ magnitude (ranging from 0 to 1) reflected the local directional consistency. By employing this weighted approach, the representative bias direction Φ_*g*_ was disproportionately determined by error vectors with larger magnitudes *w*_*i*_, thereby ensuring that the bias field primarily reflected regions of significant predictive failure.

To quantify the spatial dispersion of these biases, we simulated 5,000 trajectories, each initiated from a randomly selected valid grid cell. At each step, a virtual particle moved to the 8-neighboring cell most aligned with the direction of the local bias vector (i.e., maximizing the cosine similarity between the step direction and Φ_*g*_ ). Each trajectory was terminated if it reached a maximum of 50 steps, moved to a grid boundary, or “stalled” by remaining in the same cell for 10 consecutive steps. For each grid cell, we recorded the aggregate occupancy count (i.e., the total number of simulated steps spent in that cell across all trajectories). To isolate relative concentration of prediction biases from the total number of simulated steps, these counts were normalized over the *N*_*valid*_ valid cells to form a probability distribution *p*_*i*_ over the *N*_*valid*_ valid grid cells. The spatial entropy *H* was calculated as:

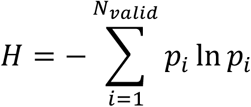

and normalized by the maximum attainable entropy ln(*N*_*valid*_) to obtain the normalized spatial entropy *H*_*norm*_ = *H* / ln *N*_*valid*_,which scaled the metric to a range of [0, 1], where 0 indicates complete concentration in a single cell and 1 indicates uniform distribution across all valid cells. This normalization ensures comparability across categories and conditions that may differ in the number of populated grid cells. A higher *H*_*norm*_ indicates that occupancy mass is broadly distributed across the Valence–Arousal space, while a lower *H*_*norm*_ indicates concentration in fewer regions, reflecting representational bias. A higher *H*_*norm*_ indicates that occupancy mass is broadly distributed across the Valence–Arousal space, while a lower *H*_*norm*_ indicates concentration in fewer regions, reflecting representational bias. The primary analysis in the main text focuses on pooled data across all categories (*N*_valid_ = *N*_*total*_), with the same procedure applied to each individual semantic category to assess category-specific bias concentration (see Supplementary Information). The error vector field, occupancy distribution, and *H*_*norm*_ were computed under two complementary cross-validation protocols. Under the LOTO protocol, all four semantic categories were represented in the training folds, and prediction errors for each category were aggregated from the held-out theme of each fold. The resulting error field characterizes the geometry of representational errors when the encoder is queried with images from categories present in training. Under the LOCO protocol, prediction errors for each category were obtained from the fold in which that entire category was held out from training. The resulting error field characterizes the geometry of representational errors when the encoder is queried with images from categories absent from training. Computing both fields with otherwise identical procedures allows differences attributable to pretraining-induced representational structure (encoder differences within a single protocol) to be distinguished from those attributable to cross-category extrapolation (within-encoder differences between protocols).

To statistically evaluate the structural properties of the error vector field, we performed directional statistical analyses. To test for a systematic ‘regression-to-the-mean’ bias, we calculated the cosine similarity between each image’s error vector **e**_*i*_ and the vector pointing from its true rating toward the geometric center of the scale (Valence = 4.0, Arousal = 4.0). Statistical significance was assessed using one-sided Wilcoxon signed-rank tests against a null hypothesis of zero alignment. Effect sizes were quantified using rank-biserial correlation (*r*_*rb*_) and Cohen’s *d*.

### Layer-Wise Probing

To investigate the network depth at which affective information is represented, we extracted [CLS] token representations from intermediate Transformer layers (layers 3, 6, 9, and 12) of each encoder. For each layer, a Ridge regression head was fitted using the same nested *α* selection procedure and the same cross-validation protocol as in the full-model analysis, with encoder weights frozen and only the source layer varied across experiments. The Ridge regularization parameter *α* was selected independently for the full-model analysis and for each layer in the layer-wise analysis through nested cross-validation, following standard practice in linear probing studies (Alain & Bengio, 2017). Each layer’s *α* was optimized for that layer’s representation, since the dimensionality and statistical properties of intermediate representations differ across depth and require layer-specific regularization. As a consequence of these differences in feature source and α selection, the layer-wise *R*^2^ and *H*_*norm*_ values are not directly comparable in absolute scale to the full-model values (which used post-projection embeddings; see Models); each set reflects the best linear decodability and error geometry achievable with the corresponding feature representation.

We performed two complementary layer-wise analyses. First, we quantified the predictive power of each layer by computing *R*^2^ on aggregated test predictions across all folds, both globally and for each semantic category. This analysis was carried out separately under LOTO and LOCO, yielding a depth-resolved profile of how much affective information each layer makes linearly decodable under within-distribution and cross-category conditions. Second, to characterize how the structure of prediction errors evolves with depth, we computed the error vector field and *H*_*norm*_ at each layer. For each layer-*l*, predicted Valence and Arousal values were obtained from the layer-*l* Ridge probe under both LOTO and LOCO, and per-image error vectors 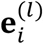 were computed and aggregated using the same grid-discretization, weighted phase-locking, and trajectory simulation procedures described above. This produced two depth profiles per encoder: 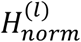 under LOTO and LOCO protocols.

The layer-wise *H*_*norm*_ profile complements the layer-wise *R*^2^ profile. *R*^2^ measures how much affective information is linearly decodable from each layer, whereas *H*_*norm*_ measures how the residual error structure that remains after linear decoding is distributed across the affective space. A layer with high *R*^2^ and low *H*_*norm*_ indicates an accurate but geometrically concentrated representation, with residual errors confined to a small number of regions. A layer with low *R*^2^ and high *H*_*norm*_ indicates a representation from which affect is not linearly decodable but whose residual errors are distributed broadly across the affective space. Together, these two profiles describe both the decodability and the geometric organization of affective information across network depth.

## Statistical Inference and Software

Model comparison (CLIP versus ViT) was assessed by paired permutation tests on fold-level *R*^2^ (LOTO: 248 folds; LOCO: 4 folds, with permutation over 248 themes as above). For LOTO we reported Cohen’s *d* for the *R*^2^ difference; for LOCO we reported *R*^2^ and *p*-values. Comparisons of *H*_*norm*_ at the full-model level were assessed by paired bootstrap tests. For each comparison, image-level error vectors were resampled with replacement (5,000 iterations) within each protocol, *H*_*norm*_ was recomputed for each resample, and the resulting distribution of paired differences was used to obtain 95% confidence intervals and two-sided *p*-values. The same procedure was applied to within-encoder comparisons of LOTO versus LOCO. For category-level paired differences, where the unit of analysis is the semantic category (n = 4), we report descriptive statistics (median and range) and exact sign tests, noting that the small number of categories limits inferential power for protocol-level comparisons. Layer-wise *H*_*norm*_ comparisons were reported descriptively in Supplementary Information. All analyses were implemented in Python using PyTorch (Paszke et al., 2019), OpenCLIP (Cherti et al., 2023), Hugging Face Transformers (Wolf et al., 2020), scikit-learn (Pedregosa et al., 2011), and Optuna (Akiba et al., 2019).

## Data Availability

The OASIS image set and normative valence and arousal ratings, are publicly available for download from osf.io/6pnd7.

## Code Availability

All custom analysis code, including the error vector field construction, the total number of trajectory steps spent in each grid cell simulation, and spatially resolved cluster analysis, is available at https://github.com/ShoheiTsu/Geometric-Affect-Analysis.git.

## Results

### Model Comparison

CLIP (ViT-B/32) and the classification-pretrained ViT (ViT-B/16) were compared under nested LOTO with a joint Ridge regression predicting Valence and Arousal (Fig. 2). Results are summarized in Table 1. Under LOTO (248 folds, 248 theme bases), CLIP achieved overall *R*^2^ = 0.68 and the ViT 0.47 (Δ*R*^2^ = 0.21, 95% CI [0.17–0.26]; paired theme-level permutation tests: *p* < 0.001). All *R*^2^ values are based on aggregated test predictions.

**Table 1.**
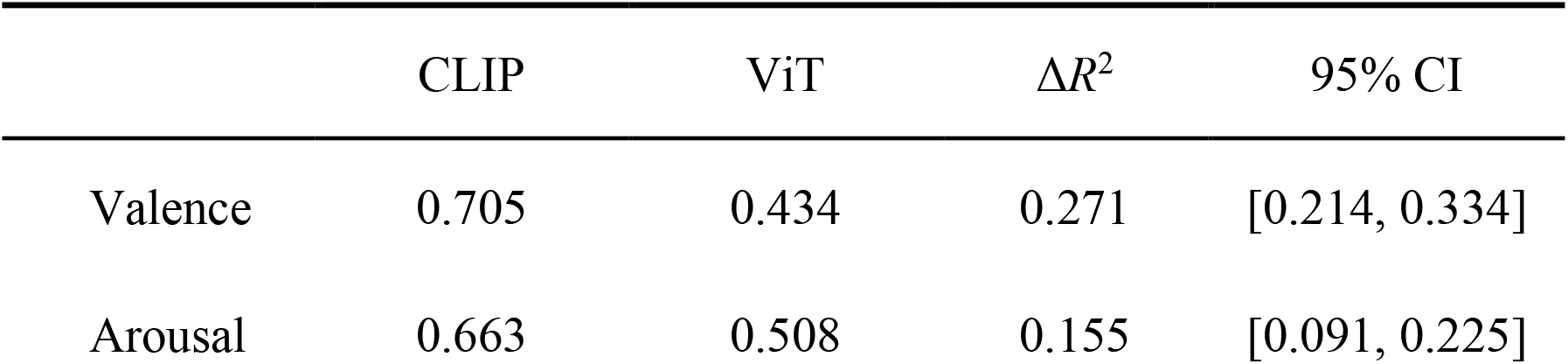
Predictive performance under Leave-One-Theme-Out (LOTO) cross-validation.

**Figure 2.**
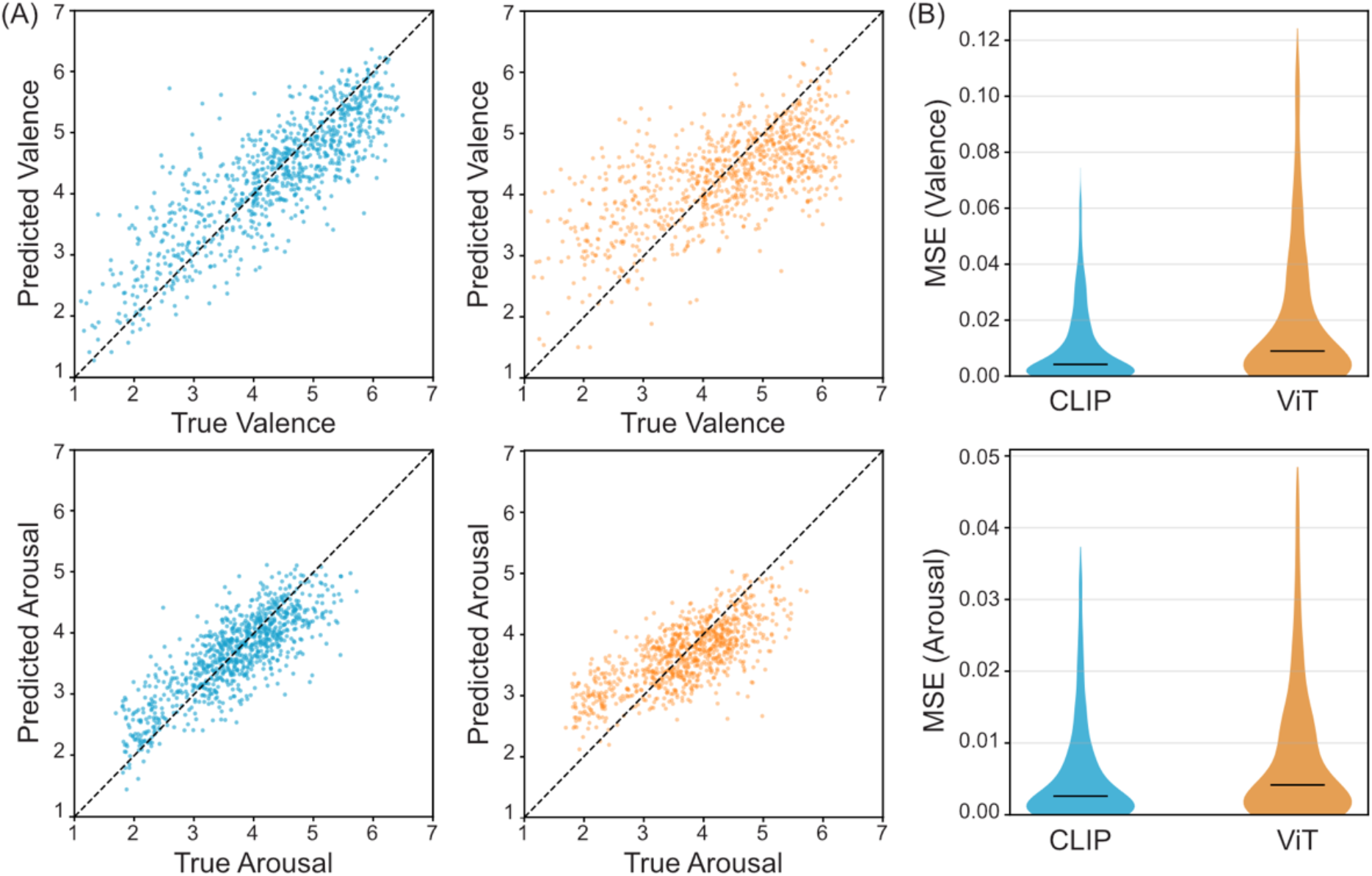
Performance comparison between CLIP and ViT under Leave-One-Theme-Out (LOTO) cross-validation. **(A) Scatter plots of true versus predicted ratings.** The top panels show results for Valence, and the bottom panels show results for Arousal. Predictions from CLIP (blue) and ViT (orange) are plotted against ground-truth ratings across 900 images. The dashed diagonal lines indicate ideal performance where predicted values equal true ratings. CLIP shows tighter clustering along the diagonal for both dimensions, indicating higher predictive accuracy and better generalization to novel semantic themes compared to the classification-pretrained ViT. **(B) Distribution of prediction errors**. The violin plots show the Mean Squared Error (MSE) distribution for Valence (top) and Arousal (bottom) across all images. Black horizontal bars indicate the median MSE. Note that MSE values were calculated after normalizing the ratings to a 0 to 1 scale. The wider and higher distribution for the ViT baseline, particularly for Valence, reflects its increased susceptibility to large predictive failures when encountering novel thematic content. In contrast, the CLIP error distribution is more concentrated near zero, demonstrating more robust representational quality for affective regression tasks.

LOCO results showed that the performance gap was strongly category-dependent (Fig. 3 and Table2). When aggregated across all four nested LOCO folds, CLIP achieved a mean *R*^2^ = 0.47, whereas the ViT fell to *R*^2^ = 0.13, yielding a CLIP–ViT gap of Δ*R*^2^ = 0.35 (95% CI [0.31–0.39]; paired permutation tests, *p* < 0.001). At the dimension-specific level, a striking dissociation emerged. CLIP maintained robustly positive *R*^2^values for both Valence and Arousal across all four categories. In contrast, while the ViT’s Valence predictions remained positive, its Arousal predictions collapsed to near-zero or negative values.

**Figure 3.**
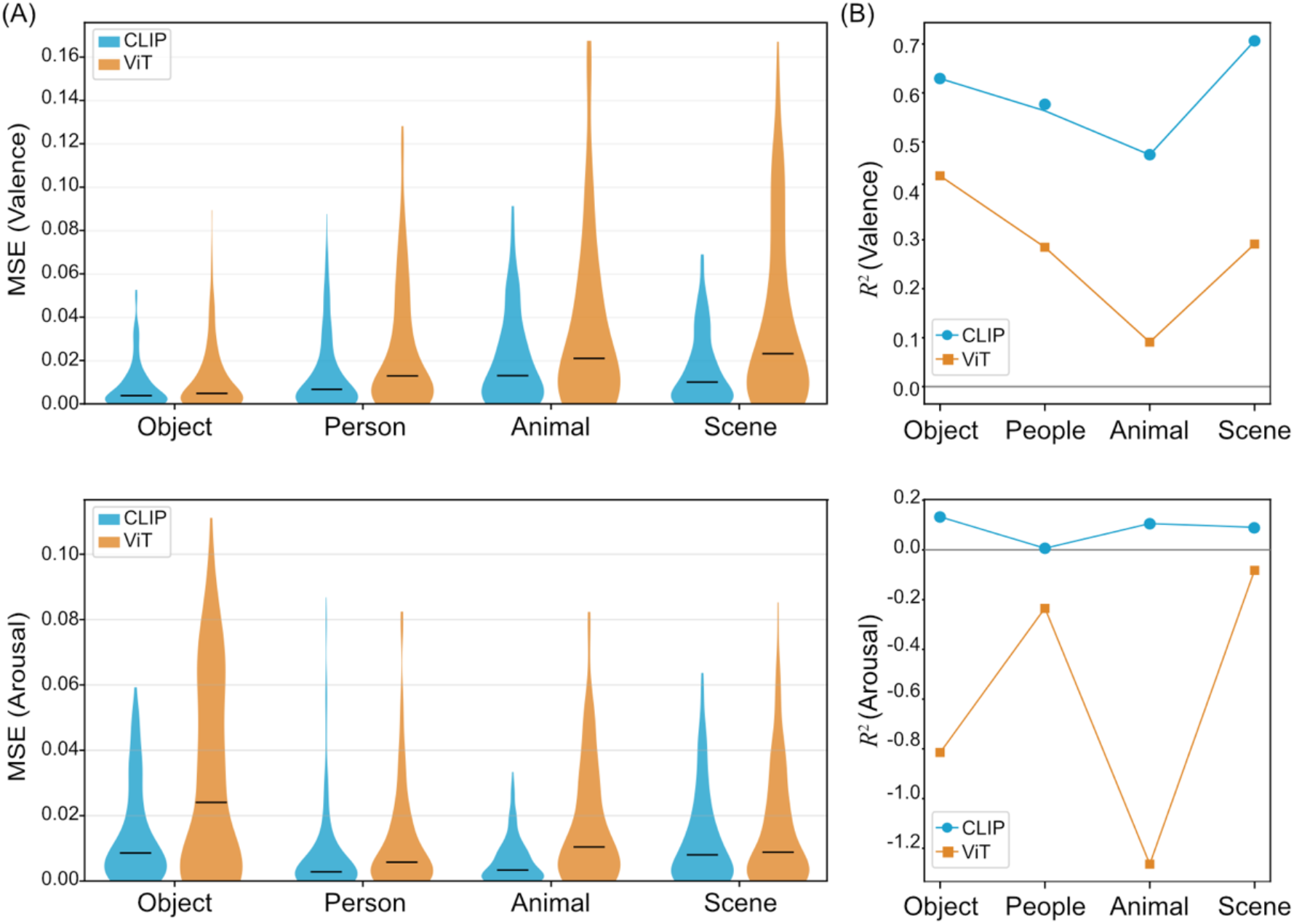
Category-specific generalization performance under Leave-One-Category-Out (LOCO) cross-validation. **(A) Distribution of prediction errors**. The violin plots illustrate the distribution of Mean Squared Error (MSE) for Valence (top) and Arousal (bottom) across the four semantic categories (Object, Person, Animal, and Scene). Each category represents a held-out test fold in the LOCO protocol. While CLIP (blue) maintains low and concentrated error distributions across all categories, the classification-pretrained ViT (orange) exhibits significantly larger errors and wider variances, reflecting a loss of robustness when encountering novel semantic classes. For both panels, error magnitude is quantified by MSE calculated on a 0 to 1 scale, with black horizontal bars indicating the median. **(B) Category-wise predictive accuracy**. The line plots show the *R*^2^ scores for each held-out category. CLIP (blue) maintains positive *R*^2^ values for both dimensions across all categories. In contrast, the ViT baseline (orange) shows a dramatic drop in performance; while its Valence predictions remain positive, its Arousal predictions collapse to negative *R*^2^ values across all categories, particularly for the Animal category. This indicates that ViT’s representations of arousal are heavily dependent on specific training categories and fail to generalize. Detailed statistical summaries, including *R*^2^, Δ*R*^2^, and 95% confidence intervals, are provided in Table 2.

### Spatially Resolved Analysis in Valence–Arousal Space

The spatially resolved analysis revealed significant CLIP-advantage clusters across the full Valence–Arousal range, including both negative and positive Valence extremes as well as intermediate Arousal levels (Fig. 4; cluster-corrected *p* < 0.05). No cluster reached significance in favor of the ViT in the overall analysis or in any individual category. Applying the same cluster-corrected analysis within each semantic category showed consistent CLIP advantage at affective extremes across all four categories, with no significant cluster observed near the neutral center in any category (Supplementary Fig S2).

**Table 2.**
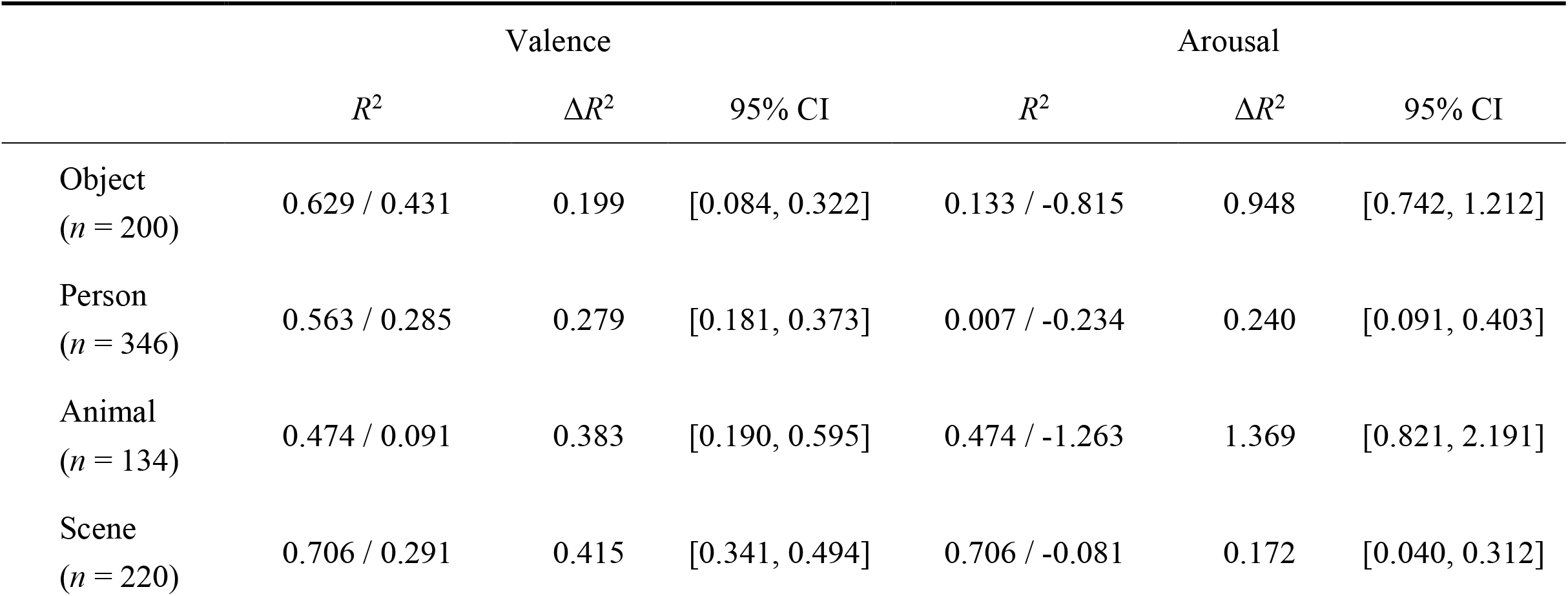
Predictive performance under Leave-One-Category-Out (LOCO) cross-validation.

|  | Valence |  |  | Arousal |  |  |
| --- | --- | --- | --- | --- | --- | --- |
| | $R^2$ | $\Delta R^2$ | 95% CI | $R^2$ | $\Delta R^2$ | 95% CI |
| Object<br>($n = 200$) | 0.629 / 0.431 | 0.199 | [0.084, 0.322] | 0.133 / -0.815 | 0.948 | [0.742, 1.212] |
| Person<br>($n = 346$) | 0.563 / 0.285 | 0.279 | [0.181, 0.373] | 0.007 / -0.234 | 0.240 | [0.091, 0.403] |
| Animal<br>($n = 134$) | 0.474 / 0.091 | 0.383 | [0.190, 0.595] | 0.474 / -1.263 | 1.369 | [0.821, 2.191] |
| Scene<br>($n = 220$) | 0.706 / 0.291 | 0.415 | [0.341, 0.494] | 0.706 / -0.081 | 0.172 | [0.040, 0.312] |

**Figure 4.**
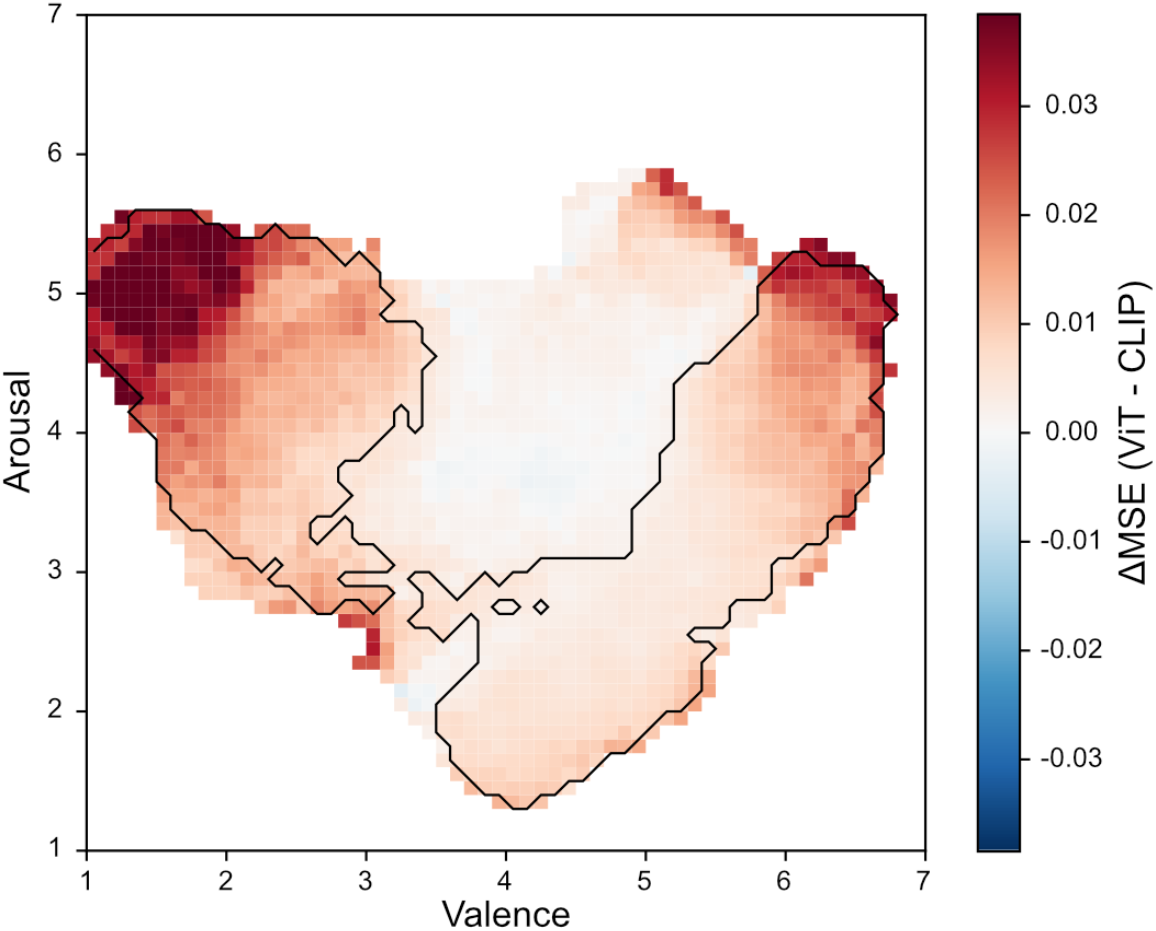
Spatially resolved performance difference in the Valence–Arousal space. The heatmap visualizes the spatial distribution of the difference in prediction error between the two encoders, defined as ΔMSE (ViT - CLIP). Red regions indicate areas where CLIP achieved lower Mean Squared Error than the classification-pretrained ViT, while blue regions would indicate a ViT advantage. Black contours enclose significant clusters where the performance gap reached statistical significance (cluster-corrected *p* < 0.05 via permutation testing). The results demonstrate a pervasive CLIP advantage across the affective space, particularly at the extremes of the Valence dimension and at intermediate to high Arousal levels. No significant clusters were observed in favor of the ViT.

### Error Vector Field and Dynamics Analysis

To characterize the directional structure and spatial concentration of prediction errors, we constructed an error vector field and simulated stochastic trajectories to generate the total number of trajectory steps spent in each grid cell (Fig. 5A). The spatial dispersion of these biases was quantified using the effective number of grid cells and normalized spatial entropy (*H*_*norm*_). When aggregated across all images under LOTO, CLIP exhibited a substantially broader distribution of prediction biases than the ViT (*H*_*norm*_: 0.61 vs. 0.55). This indicates that while CLIP’s errors are distributed across a wide range of the Valence– Arousal space, the ViT’s errors are highly concentrated within a few specific “sinks.” The divergence was most pronounced in the Person category, where CLIP’s bias was more dispersed than the ViT’s ( *H*_*norm*_ : 0.69 vs. 0.47). In contrast, the differences were markedly smaller for the Object and Scene categories (Table S2). Qualitative inspection of the error vector field showed a rotational structure in CLIP’s predictions for Person images, which contributes to the high *H*_*norm*_ value observed in this category (see Supplementary Section 3.4 for further interpretation).

**Figure 5.**
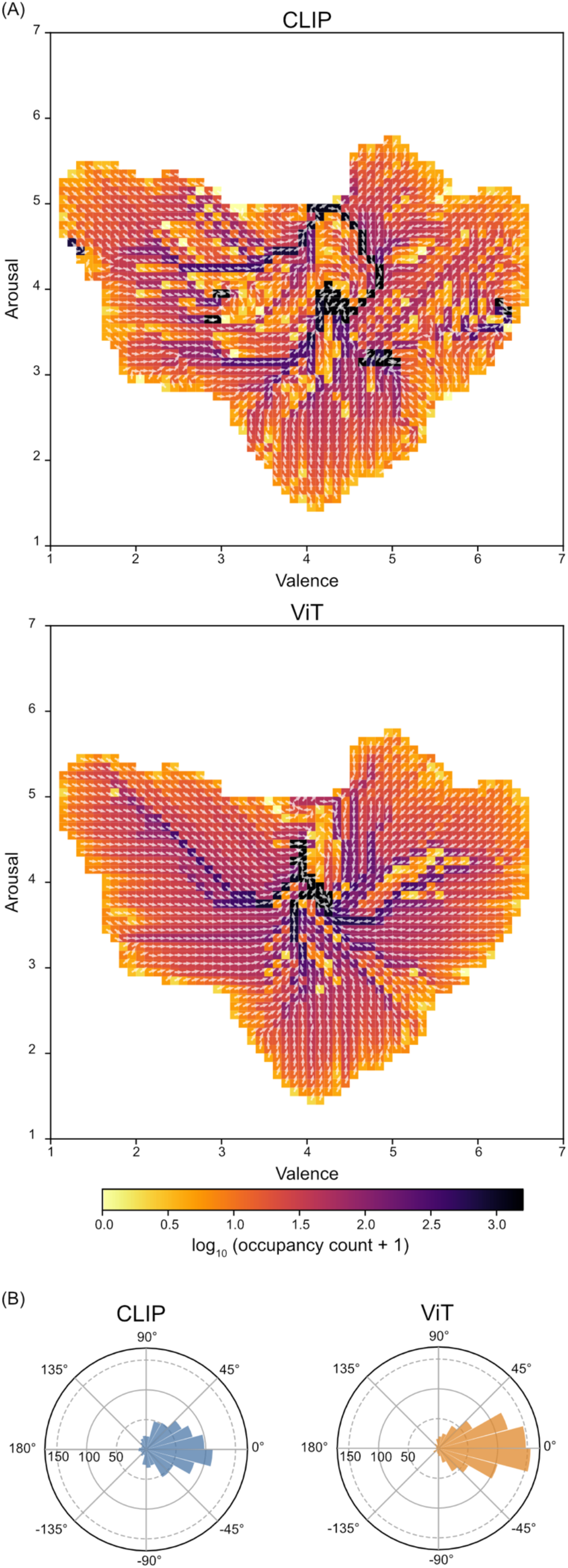
Geometric structure and stochastic dynamics of prediction biases. **(A) Trajectory occupancy maps and error vector fields**. The heatmaps visualize the concentration of prediction biases for CLIP (left) and ViT (right) across the Valence– Arousal space. The color scale represents the log-transformed cumulative visit counts from a stochastic ball simulation, where movement is governed by the local error vector field (white arrows). High-occupancy regions (darker areas) indicate “bias sinks” where predictive failures consistently aggregate. The ViT exhibits a more concentrated occupancy pattern compared to CLIP, particularly showing a more severe collapse into specific affective subspaces. This spatial dispersion was quantified using effective number of grid cells and normalized entropy (*H*_*norm*_), with CLIP demonstrating higher dynamic diversity. **(B) Center-ward bias and regression-to-the-mean**. The rose diagrams illustrate the distribution of phase differences between the local error vectors and the direction toward the geometric center of the affective space (Valence = 4.0, Arousal = 4.0). A phase difference of 0° indicates an error vector pointing directly toward the center. Both models exhibit a marked center-ward bias, consistent with a regression-to-the-mean effect. However, the distribution for the ViT (orange) is more tightly concentrated around 0° compared to CLIP (blue), indicating a more rigid and pervasive representational collapse toward the neutral center.

To investigate the structural bias of prediction errors, we analyzed the alignment of error vectors toward the geometric center (Valence = 4.0, Arousal = 4.0). Circular distributions of the phase difference ( *δ* ) between error vectors and the centerward direction revealed a marked concentration near 0° for both models, confirming a pervasive regression-to-the-mean tendency (Fig. 5B). This center-oriented bias was further quantified using cosine similarity (cos *δ*). While both encoders exhibited a strong center-ward pull (CLIP: *median* = 0.79, *d* ≈ 0.91; ViT: *median* = 0.91, *d* ≈ 1.62) paired comparisons indicated that the alignment toward the center was stronger in ViT than in CLIP (*median* Δcos = 0.056, *p* < 0.001, *r*_*rb*_(Δ) ≈ 0.36). While both models regress toward the mean, the higher cosine similarity and effect sizes in ViT indicate a more severe representational collapse toward the center, whereas CLIP’s errors maintain a higher degree of directional diversity. Category-level analyses further confirmed that while center-ward bias is a pervasive feature of both models, its intensity is modulated by semantic content, with the gap between encoders being most pronounced in the Object category (see Supplementary Material Section 4).

### Layer-wise Representation of Affective Information

To investigate whether affective information is localized to specific layers or distributed across the network depth, we performed layer-wise linear probing (layers 3, 6, 9, and 12; Fig. 6). For both encoders, single-layer probes yielded substantially lower *R*^2^ values compared to the full model, indicating that robust affect prediction relies on the integration of features across multiple hierarchical levels rather than any single representational stage. The two encoders exhibited diverging trends across depth. CLIP showed small positive *R*^2^ values at shallow layers (layers 3-6), which decreased toward near-zero or slightly negative values at deeper layers. In contrast, the ViT exhibited positive *R*^2^ at every evaluated layer, with predictive performance increasing toward the final layer (layer 12). Furthermore, CLIP maintained a more uniform category profile across layers, whereas the ViT showed increasing cross-category dispersion as depth increased, peaking at the deepest layer (Supplementary Fig. S7).

**Figure 6.**
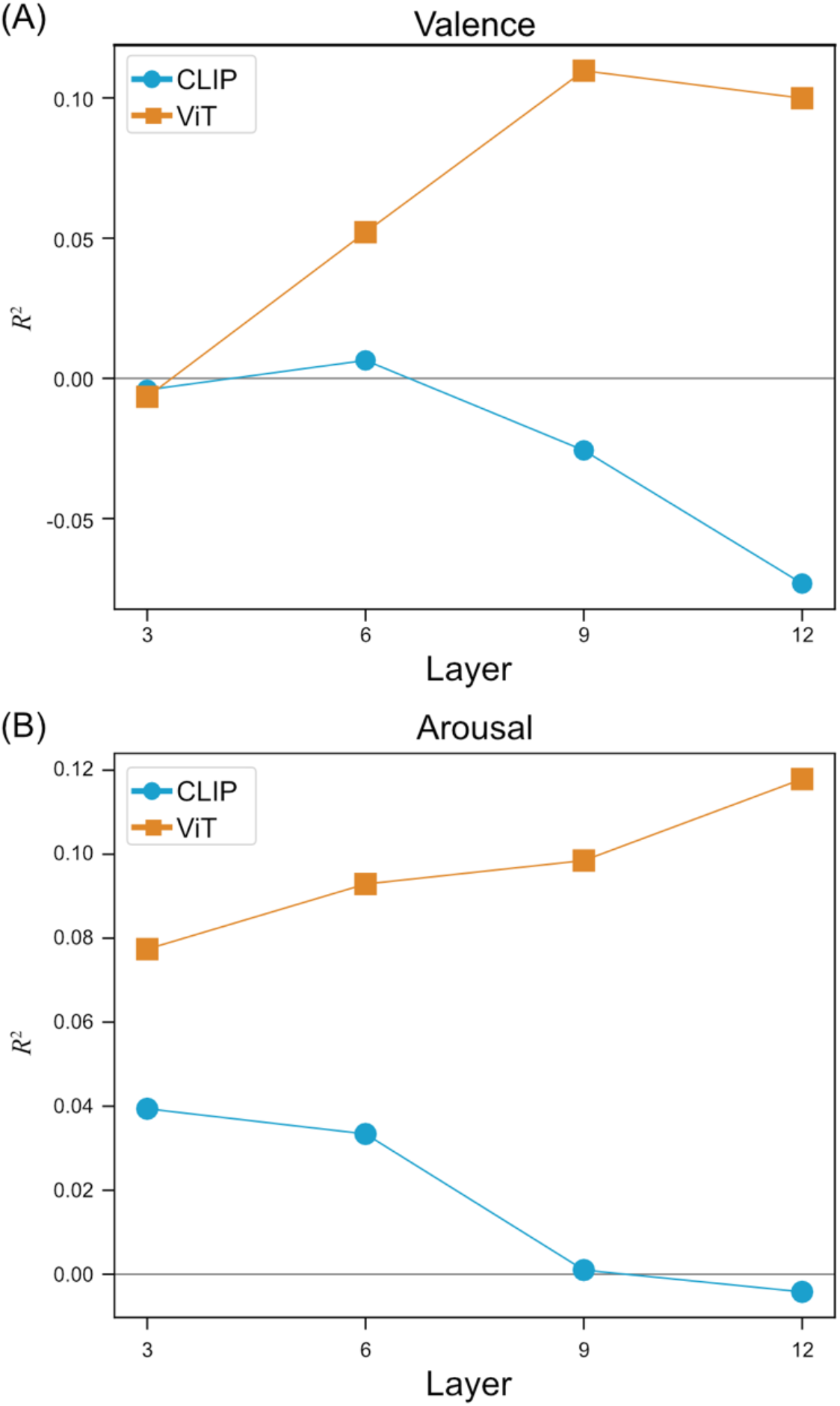
Layer-wise linear probing across network depth. **(A) Predictive performance for Valence by layer**. The line plots indicate the *R*^2^ scores obtained by fitting a Ridge regression head to intermediate features from layers 3, 6, 9, and 12 under Leave-One-Theme-Out (LOTO) cross-validation. While CLIP (blue) shows minimal and decreasing predictive power at deeper layers, the classification-pretrained ViT (orange) exhibits a steady increase in performance toward the final layer. **(B) Predictive performance for Arousal by layer**. Similar to Valence, the ViT shows increasing *R*^2^ values as the network depth increases. In contrast, CLIP exhibits low performance that further diminishes in the deepest layers. For both panels, the substantially lower *R*^2^ values compared to the full model (which utilizes the final embedding) suggest that robust affect prediction requires the integration of high-level features represented at the network’s output. The diverging trends indicate that while CLIP distributes affective semantics across the network hierarchy, the ViT increasingly concentrates predictive, yet category-bound, visual features toward its final layers.

To examine the geometric structure of layer-wise prediction errors, we computed *H*_*norm*_ at each layer under both LOTO and LOCO protocols. The layer-wise *H*_*norm*_ values used [CLS] token features from intermediate layers and layer-specific Ridge regularization, and are therefore not directly comparable in absolute scale to the full-model *H*_*norm*_ values reported in Supplementary Table S2. Neither encoder exhibited a monotonic depth trend in pooled *H*_*norm*_ under either protocol, and category-resolved profiles showed substantial heterogeneity across the combinations of category, protocol, and depth. Detailed layer-wise *H*_*norm*_ results, including pooled and per-category values, are presented in Supplementary Information (Supplementary Fig. S8 and S9).

## Discussion

### Cross-Category Generalization and the Limits of Random-Split Evaluation

Our results demonstrate that CLIP consistently outperforms the classification-pretrained ViT in affective image prediction, with the performance gap most pronounced when the models are required to generalize to unseen semantic categories. By holding the backbone family constant and selecting the stronger ImageNet-trained variant as the classification baseline, the comparative design ensures that observed differences can be attributed primarily to the pretraining objective rather than to architectural confounders. Although certain architectural parameters varied between the primary models (CLIP-B/32 vs. ViT-B/16), supplementary analyses confirm that the pretraining objective remains the dominant determinant of representational structure regardless of these variations.

Cross-category generalization has been an implicit but rarely tested requirement of affective image analysis systems. Standardized affective image collections such as IAPS (Lang, 1995), NAPS (Marchewka et al., 2014), and OASIS (Kurdi et al., 2017) are organized around heterogeneous semantic categories (animals, faces, scenes, objects, and social situations), yet most computational evaluations on these datasets rely on random-split cross-validation, which mixes categories in training and test folds. Model performance reported in prior work therefore largely reflects within-category interpolation rather than cross-category extrapolation, leaving the question of categorical robustness unaddressed. The LOCO protocol adopted here directly tests this property by withholding entire semantic categories at test time, revealing a vulnerability that random-split protocols systematically conceal: classification-pretrained features that perform competitively under random splits show diminished predictive power when forced to generalize across category boundaries.

The LOCO results revealed a dissociation in the classification-pretrained ViT’s generalization capacity. While the model retained modest predictive power for Valence across categories, its ability to predict Arousal collapsed to near-zero or negative levels when categorical boundaries were crossed, particularly for the Object and Animal categories. This selective collapse aligns with theoretical accounts of the Valence and Arousal dimensions as partially independent components of affective experience, with distinct semantic and perceptual determinants (Russell, 1980). Valence judgments are typically driven by category-level semantic content (whether a depicted entity is broadly pleasant or aversive), which can be partially preserved by classification-trained features encoding entity identity. Arousal depends more on contextual and relational properties such as threat, novelty, and social dynamics, which are not captured by object-level visual statistics (Lang, 1995; Marchewka et al., 2014). Neuroimaging evidence further indicates that arousal processing recruits distributed networks integrating interoceptive and contextual information rather than localized object-recognition pathways (Barrett & Simmons, 2015). ImageNet-trained features, optimized for entity discrimination, can therefore support residual Valence prediction across categories but provide little signal for Arousal. The fact that this dissociation appears specifically in the LOCO protocol, and not in LOTO, indicates that it is a property of category-level extrapolation rather than a general failure of arousal prediction.

CLIP’s language-aligned representations supported generalization across all four semantic categories, consistent with prior arguments that affect is more directly tied to semantic content than to low-level-visual structure (Borth et al., 2013; Machajdik & Hanbury, 2010). Within affective image analysis, this principle has been articulated through frameworks such as adjective–noun pairs (Borth et al., 2013) and feature systems inspired by psychology and art theory (Machajdik & Hanbury, 2010). Language–image contrastive training operationalizes a similar bridge at scale, aligning image embeddings with the textual descriptions humans use to describe scenes, including affectively loaded vocabulary such as threatening, soothing, joyful, and disturbing. Psychological constructionist theories of emotion hold that affective experience arises when core affect is conceptualized through learned, language-supported emotion concepts rather than read off from invariant perceptual features (Barrett & Simmons, 2015; Lindquist et al., 2006). Under this view, language alignment is not merely a useful auxiliary signal but a natural way to introduce the conceptual structure that affective categorization requires. ImageNet classification, by contrast, optimizes for object identity and is largely indifferent to the relational semantics that determine affective significance, providing a structural account of its failure under categorical extrapolation.

These findings extend prior computational work on the OASIS dataset. Earlier approaches based on global image properties achieved Valence prediction in the modest range previously reported (Redies et al., 2020), and a CNN-based study reported low Valence performance under random-split evaluation (Priyadarshani & Miyapuram, 2025). Transformer-based encoders had not been applied to OASIS, and generalization across entire semantic categories had not been evaluated. The improvement obtained by CLIP under the more stringent LOTO protocol therefore represents a substantive shift in what is achievable on this benchmark, and the persistence of CLIP’s advantage under LOCO, where the classification-pretrained ViT diminishes to negligible performance, confirms that the gain is not an artifact of training-test similarity. The supplementary three-model comparison, which includes ImageNet-ViT-B/32 as a reference, confirms that CLIP’s advantage over the classification baseline is not attributable to patch resolution or backbone capacity but persists when architecture is matched exactly.

### Geometric Structure of Prediction Errors

The spatially resolved error analysis localized CLIP’s advantage to specific regions of the Valence–Arousal plane. Significant CLIP-advantage clusters spanned both negative and positive Valence extremes as well as intermediate-to-high Arousal levels, indicating that the language-aligned representation is most beneficial where content carries strong and unambiguous emotional significance (Lang, 1995; Marchewka et al., 2014). Near the neutral center of the affective space, the two encoders did not differ significantly, consistent with the view that affectively neutral images are not fully accounted for by semantic content alone and may depend more on perceptual factors and individual response variability (Barrett & Simmons, 2015; Lindquist et al., 2006; Zhao et al., 2022). The pattern held across all four categories: CLIP-advantage clusters appeared at affective extremes in every category, and no category yielded a ViT-advantage cluster. In the high-Arousal, moderately positive Valence region of the Scene category, no significant cluster was observed for either encoder, a null result consistent with the view that affective responses to scenic content in this region depend on perceptual properties such as luminance, openness, and spatial structure (Berto, 2014; Kaplan & Kaplan, 1989) that neither encoder is specifically optimized to capture. Spatial difference mapping between LOTO and LOCO predictions further showed that the regions of greatest LOCO-induced error increase coincided across encoders for Object images but diverged for Animal, Person, and Scene (Supplementary S2.4), indicating that LOCO-induced errors in the latter three categories reflect encoder-specific representational gaps rather than a shared difficulty inherent to specific regions of the affective space.

The geometric analysis of prediction errors, complemented by trajectory-based occupancy simulations, offers a structural interpretation of these performance gaps. Both encoders exhibited a center-oriented regression bias, but the spatial dispersion of their prediction errors differed systematically. Under within-distribution evaluation (LOTO), the language-aligned encoder showed consistently higher normalized spatial entropy (*H*_*norm*_) of the error distribution than the classification-pretrained ViT across all four semantic categories, indicating a broader error topology when category coverage is preserved in training. The classification-pretrained ViT, by contrast, exhibited lower *H*_*norm*_ values, with prediction errors more concentrated within a limited number of high-error regions of the affective space, structurally reminiscent of the representational compression observed in supervised classification training (Papyan et al., 2020). This concentrated error topology may render the classification-pretrained model more susceptible to shortcut learning (Geirhos et al., 2020), where predictions rely on spurious correlations that fail to transfer across categorical boundaries.

Comparing LOTO and LOCO error geometries reveals an asymmetric pattern that refines the interpretation of these representational differences. Under cross-category evaluation (LOCO), the *H*_*norm*_ advantage of the language-aligned encoder observed under LOTO was substantially reduced, with *H*_*norm*_ values converging toward intermediate ranges for both encoders. The language-aligned encoder showed larger reductions in *H*_*norm*_ under LOCO than the classification-pretrained encoder, particularly for the Person and Animal categories, while the classification-pretrained encoder showed smaller and more uniform changes across categories. This indicates that the broader error topology of language-aligned representations is most evident under within-distribution conditions, while category-level extrapolation introduces additional concentration of errors that affects both encoders. The aggregate prediction accuracy advantage of the language-aligned encoder under LOCO (substantial improvements in *R*^2^ across categories) thus does not translate into a proportional advantage in the spatial dispersion of residual errors, suggesting that *R*^2^ and *H*_*norm*_ capture distinct aspects of representational quality: the former measures how much affective signal is recovered, while the latter measures how the residual error is distributed across the affective space.

### Layer-wise Dynamics of Affective Representations

Layer-wise probing revealed how affective information is organized across network depth in the two encoders. The classification-pretrained ViT exhibited progressively higher single-layer *R*^2^ toward the final layer under LOTO, with *R*^2^ increasing across depth. CLIP’s intermediate layers, by contrast, yielded persistently weak single-layer *R*^2^, often dropping below zero at greater depths. The local superiority of the classification-pretrained model in single-layer *R*^2^ did not translate into superior cross-category generalization. Under LOCO, the deeper layers of the ViT yielded strongly negative *R*^2^ values for the Object and Animal categories. This selective collapse indicates that classification-based pretraining accumulates category-biased visual regularities layer by layer, reflecting the texture-based biases typical of ImageNet-trained models (Geirhos et al., 2019). These features provide superficial predictive power within known domains but fail to generalize across category boundaries.

The persistent weakness of linear decoding from CLIP’s intermediate layers indicates that language-aligned pretraining distributes affective semantics across the network hierarchy rather than localizing them in any single layer (Park & Kim, 2022). The pretraining objective thus determines how affective information is organized across depth: the classification-pretrained ViT concentrates predictive but category-bound features in its later layers, whereas CLIP distributes affective semantics across depth at the cost of intermediate-layer linear decodability.

We additionally examined the spatial dispersion of layer-wise prediction errors using *H*_*norm*_ at four intermediate Transformer layers (Supplementary Figs. S7 and S8). Neither encoder exhibited a monotonic depth trend in *H*_*norm*_, and category-resolved profiles showed neither a consistent encoder ordering nor a consistent depth trend within any of the four semantic categories under either protocol. The layer-wise *H*_*norm*_ values are not directly comparable in absolute scale to the full-model *H*_*norm*_ values reported in the Results, since the layer-wise probes use [CLS] token features from intermediate layers rather than the encoder’s final image embeddings, and use layer-specific Ridge regularization. The full-model *H*_*norm*_ comparisons therefore provide the principal evidence for the encoder-level difference in error topology, while the layer-wise profiles indicate that this difference is not simply propagated through intermediate layers but is mediated by category- and protocol-specific factors.

### Implications for Representation Learning and Affective Computing

The pretraining objective of a frozen vision encoder has direct consequences for the reliability of affective computing systems deployed in real-world settings. In application contexts where the operational image distribution overlaps closely with the training distribution, such as scene-focused environmental affect estimation, classification-pretrained encoders remain a computationally efficient choice and yielded positive prediction performance for Scene images in our experiments. Applications that must operate over heterogeneous content, including affect-aware content recommendation, open-domain stimulus selection for psychological research, affective dialogue and tutoring systems, and emotion-sensitive human–computer interaction, necessarily encounter semantic categories absent from any fixed training distribution. The decline in classification-pretrained ViT performance under the LOCO protocol indicates that such systems risk overestimating their own reliability when deployed beyond their training domain, performing competitively on familiar content but failing unpredictably on novel categories. Language-aligned encoders mitigate this categorical vulnerability without requiring affect-specific fine-tuning, providing a more reliable frozen feature extractor for open-domain affective computing.

Beyond the affective computing domain, our findings contribute to the broader literature on representation learning by showing that pretraining objectives shape not only aggregate downstream performance but also the geometric organization of learned representations and their robustness to category-level distribution shifts. The combination of cross-category evaluation (LOTO vs. LOCO) and geometric analysis (*H*_*norm*_ of error fields across protocols) provides a methodological template for diagnosing categorical brittleness in frozen encoders that is applicable beyond affective tasks. We suggest that LOCO-style cross-category evaluation be incorporated into the standard validation pipeline for affective image analysis systems prior to deployment, as it exposes a form of categorical brittleness that random-split evaluation cannot detect.

### Limitations

Several limitations of the current design merit consideration. First, the comparison covers only two pretraining objectives within a single backbone family. Including self-supervised methods such as DINOv2 (Oquab et al., 2024) and masked autoencoders (He et al., 2022) would situate language supervision within a broader landscape of pretraining strategies. Second, all analyses are based on a single dataset. Replication on other standardized affective image collections such as IAPS (Lang, 1995) and NAPS (Marchewka et al., 2014), as well as on larger and more heterogeneous web-scraped collections, will be important for establishing the generality of the present findings. Third, we used frozen encoder features throughout. Task-specific fine-tuning on affective labels may benefit different pretraining objectives differentially, and the interaction between pretraining objective and fine-tuning depth warrants systematic investigation. Finally, OASIS ratings reflect affective responses to static images viewed in isolation, and how the present findings extend to dynamic content, contextual framing, multimodal affective signals, and individual differences in affective response remains an open question for future work.

## Conclusions

We compared two Vision Transformer encoders, CLIP (ViT-B/32) and an ImageNet-21k–pretrained ViT (ViT-B/16), as frozen feature extractors for continuous Valence– Arousal prediction on the OASIS dataset, holding the backbone family constant to isolate the contribution of the pretraining objective. Three convergent findings emerge from this comparison. First, CLIP consistently outperforms the classification-pretrained ViT across both theme-level and category-level generalization, with the gap widening sharply under Leave-One-Category-Out evaluation, where classification-pretrained features fail to support Arousal prediction across novel semantic categories. Second, a geometric analysis of prediction errors using normalized spatial entropy r reveals that the two representations differ in the spatial dispersion of their failures. Under LOTO evaluation, language-aligned features distribute errors more broadly across the affective space than classification-pretrained features across all categories. While both encoders exhibit more concentrated error distributions under LOCO, the classification-pretrained ViT undergoes a more severe representational collapse, whereas CLIP maintains a relatively more distributed error topology. Third, layer-wise probing reveals that affective information is distributed across network depth in CLIP but concentrated in deeper, category-bound layers of the classification-pretrained ViT, mirroring the texture-bias and category-anchored statistics characteristic of ImageNet-trained representations. The layer-wise effective dimensionality profile further shows that the classification-pretrained ViT progressively compresses its residual error structure across depth—a structural signature of categorical brittleness—whereas CLIP maintains a more stable error geometry. These findings suggest that the pretraining objective, rather than architectural capacity or data scale alone, is a primary determinant of whether a learned representation captures the semantic structure that supports affective prediction across categories. For affective computing systems intended to operate over heterogeneous content, including content recommendation, stimulus selection for psychological research, and emotion-sensitive human–computer interaction, language-aligned encoders provide a more reliable frozen feature extractor than classification-pretrained alternatives. We propose that LOCO evaluation be adopted as a standard diagnostic for affective image analysis systems, since it exposes a form of categorical brittleness that random-split evaluation systematically conceals. Future work extending this comparison to self-supervised pretraining objectives, additional standardized affective image datasets, and the interaction between pretraining objective and fine-tuning depth will further situate language supervision within the broader landscape of representation learning.

## Availability of data and material

The datasets used and/or analyzed during the current study are available from the corresponding author on reasonable request.

## CRediT authorship contribution statement

**Shohei Tsuchimoto**: Conceptualization, Data curation, Formal analysis, Investigation, Methodology, Resources, Software, Validation, Visualization, Writing - Original Draft, Writing - Review & Editing, Project administration, Funding acquisition. **Yuka O Okazaki**: Visualization, Validation, Writing - Review & Editing. **Kenichi Yuasa**: Validation, Writing - Review & Editing. **Sakura Nishijima**: Writing - Review & Editing. **Mebuki Izumiya**: Writing - Review & Editing. **Makoto Hagihara**: Writing - Review & Editing. **Ryo Fujihira**: Visualization, Validation, Writing - Review & Editing. **Keiichi Kitajo**: Writing - Review & Editing, Supervision, Funding acquisition.

## Acknowledgements

This work was supported by KAKENHI (JP21K13758, JP22KJ3102 and JP26K16952) from the Japan Society for the Promotion of Science (JSPS), and JST Moonshot R&D (JPMJMS2292-1-03).

