## Supplementary Information for "Pretraining Objective Shapes Cross-Category Generalization in Affective Image Prediction: A Geometric Comparison of Vision Transformer Encoders"

#### 1. Preliminary Model Selection and Comparison

*1.1. Selection of Classification-Pretrained Baseline (ViT-B/16 vs. ViT-B/32)*

To ensure a rigorous comparison between language-aligned and classification-pretrained encoders, we first evaluated which variant of the ImageNet-21k-pretrained Vision Transformer (ViT-B) served as the strongest baseline. Although CLIP uses a patch size of 32 (ViT-B/32), preliminary tests demonstrated that the classification-pretrained ViT-B/16 consistently outperformed the ViT-B/32 variant across all evaluation metrics. Specifically, ViT-B/16 yielded higher mean $R^2$ scores under both LOTO and LOCO cross-validation protocols. Consequently, we selected ViT-B/16 as the primary baseline to provide the most challenging comparison for CLIP. This ensures that any observed performance advantages for CLIP are attributable to its pretraining objective rather than to architectural superiorities in patch resolution (Fig. S1 and Table S1).

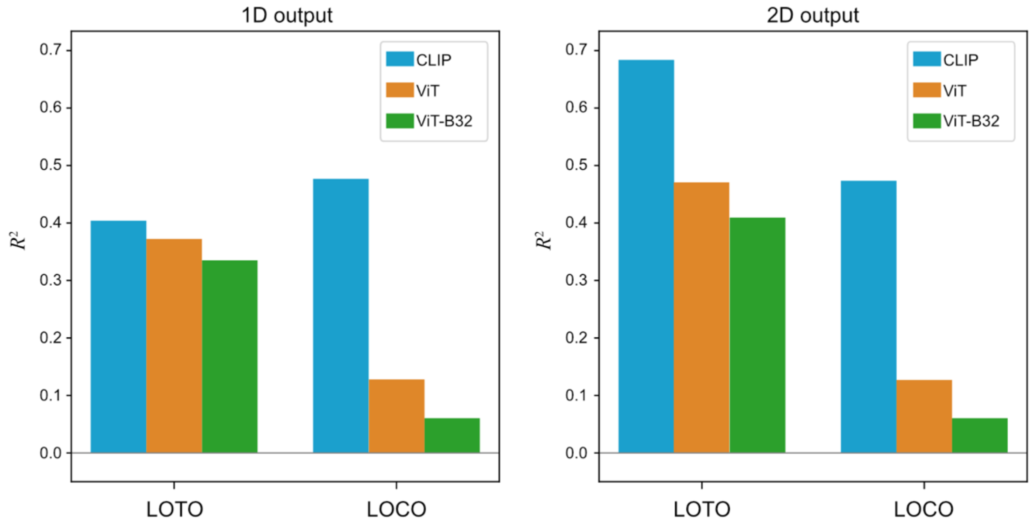


**Figure S1. Comparative performance across encoder variants and prediction strategies.** The bar charts illustrate the mean $R^2$ scores under LOTO and LOCO protocols for CLIP (ViT-B/32) and two classification-pretrained ViT variants (ViT-B/16 and ViT-B/32). **(Left)** Performance under independent 1D output prediction for Valence and Arousal. **(Right)** Performance under joint 2D output prediction.

**Table S1. Preliminary performance comparison ($R^2$) across encoder variants and prediction strategies under LOTO cross-validation.**

| | $R^2$ (1D) | | $R^2$ (2D) | |
|---|---|---|---|---|
| | Valence | Arousal | Valence | Arousal |
| CLIP (ViT-B32) | 0.376 | 0.4322 | 0.705 | 0.663 |
| ViT (ViT-B/16) | 0.295 | 0.449 | 0.434 | 0.508 |
| ViT (ViT-B/32) | 0.280 | 0.390 | 0.394 | 0.425 |

*1.2. Comparative Analysis of Prediction Strategies (1D vs. 2D Output)*

We further investigated whether predicting Valence and Arousal dimensions independently (1D output) or jointly (2D output) provided better representational stability. In the 2D output configuration, the Ridge regression $\alpha$ parameter was optimized for the joint prediction space via nested cross-validation. As shown in the supplementary figure, the 2D joint prediction strategy yielded consistently higher $R^2$ values compared to the independent 1D strategy for both encoders. This improvement suggests that the joint modeling of affective dimensions better captures the inherent correlations between Valence and Arousal, leading to more robust generalization. Based on these results, the 2D joint prediction framework was adopted for all primary analyses in the main text.

**2. Spatially Resolved Performance Analysis within Semantic Categories.**

*2.1. Spatial Mapping Procedure*

To identify regions of the Valence–Arousal space where the predictive performance of the two encoders diverged within specific semantic domains, we performed a spatially resolved error analysis for each category (Object, Person, Animal, and Scene). We utilized predictions from the Leave-One-Theme-Out (LOTO) protocol to ensure maximum coverage of the affective space, as LOTO yields a test prediction for every image in the dataset. A regular grid was superimposed on the Valence–Arousal map with a step size of 0.1 units. At each grid intersection, a local neighborhood was defined by a radius of 0.5 units. Within each neighborhood containing at least five images, we performed a paired Wilcoxon signed-rank test to compare $MSE_{CLIP}$ and $MSE_{ViT}$. The local effect size, $\Delta$MSE, was defined as the mean difference ($MSE_{ViT}$ - $MSE_{CLIP}$), where positive values indicate a performance advantage for CLIP.

*2.2. Statistical Significance and Cluster Correction*

Multiple comparisons across the spatial grid were controlled using a permutation-based cluster test with 5,000 iterations. For each iteration, the signs of the error differences were randomly flipped at the image level, and local statistics were recomputed. Contiguous cells exceeding an uncorrected threshold of $p < 0.05$ (based on 4-connectivity) were grouped into clusters. A null distribution was constructed from the maximum cluster sizes observed across all permutations, providing a corrected $p$-value for the clusters observed in the original data.

*2.3. Results of Category-Specific Mapping*

As illustrated in Figure S2, the category-specific analysis revealed significant CLIP-advantage clusters across all four semantic categories. These clusters were primarily localized at the affective extremes of the Valence dimension. Notably, no significant clusters favored the ViT baseline in any category. The spatial distribution of these clusters remained consistent across different semantic contents, suggesting that CLIP's superior representational quality at affective extremes is a robust property that transcends categorical boundaries. This finding further supports the conclusion that language-aligned pretraining facilitates a more semantically grounded and generalized mapping of human affect than traditional classification-based objectives.

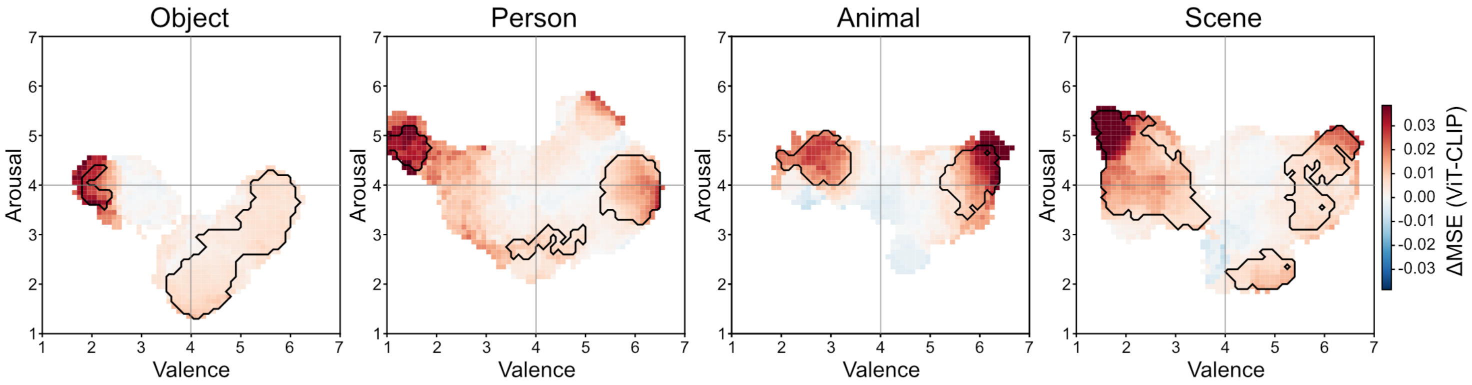


**Figure S2. Category-specific spatial performance differences in the Valence–Arousal space.** The heatmaps visualize the spatial distribution of ΔMSE (ViT - CLIP) for each individual semantic category (Object, Person, Animal, and Scene) based on the LOTO cross-validation data. Red regions indicate areas where CLIP achieved lower prediction error than the classification-pretrained ViT baseline. Black contours enclose significant clusters where the CLIP advantage reached statistical significance (cluster-corrected $p < 0.05$ via permutation testing). Consistent with the aggregate analysis shown in Figure 4, significant CLIP advantages are predominantly localized at affective extremes (e.g., very high or low Valence) across all categories. In contrast, no significant differences were observed near the neutral center (Valence ≈ 4.0, Arousal ≈ 4.0) for any category.

*2.4. Spatial Difference Maps Between LOTO and LOCO Predictions*

To determine whether LOCO-induced performance drops were localized to specific regions of the affective space, and whether such regions were shared between encoders or were encoder-specific, we computed per-image squared error differences (LOCO − LOTO). We applied the same spatial mapping and cluster-corrected significance testing procedures as in the within-category analysis (described above for the cluster-corrected mapping in Statistical Significance and Cluster Correction and Results of Category-Specific Mapping; Fig.S3). For each encoder and semantic category, we identified Valence–Arousal grid cells where the LOCO-induced error increase exceeded chance under permutation testing (*n_perm* = 1,000, *p_uncorr* = 0.05, *cluster_alpha* = 0.05).

Significant clusters of LOCO-induced error increase were identified in all four semantic categories for both encoders. For Object images, the peak of LOCO-induced error increase fell in nearly identical regions for the two encoders (CLIP peak at V = 4.15, A = 1.35; ViT peak at V = 4.45, A = 1.45), indicating a shared region of cross-category extrapolation difficulty in low-arousal, neutral-valence content. For the remaining categories, the peak locations diverged substantially between encoders. The largest divergence occurred in the Animal category, where the CLIP peak (V = 2.65, A = 3.75) and the ViT peak (V = 5.65, A = 4.65) were separated by approximately three valence units. The Person and Scene categories showed peak locations separated by approximately one valence unit between encoders. The shared peak in the Object category suggests that certain V-A regions are difficult to predict under cross-category extrapolation regardless of the pretraining objective, consistent with images in those regions carrying ambiguous or context-dependent affective signals. The divergent peaks in the Animal, Person, and Scene categories indicate that the remaining LOCO-induced errors cannot be attributed to image difficulty alone but reflect encoder-specific representational gaps. CLIP and the classification-pretrained ViT thus fail in different regions of the affective space when generalizing to held-out semantic categories, consistent with the broader finding that pretraining objective shapes the geometry of cross-category generalization errors.

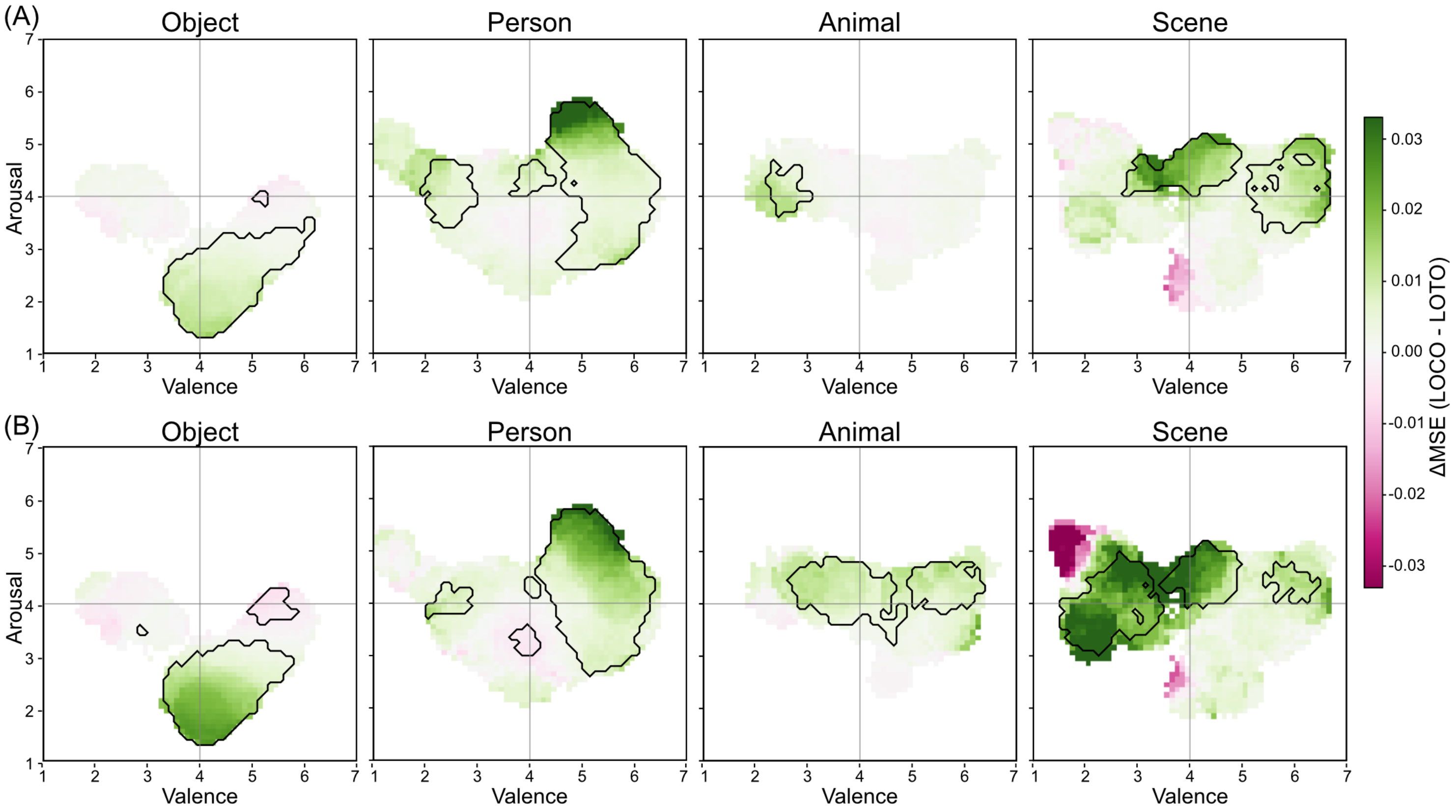


**Figure S3. Spatial difference maps of LOCO-induced prediction error increases.** The maps show the spatial distribution of squared prediction error differences (LOCO minus LOTO) across the Valence–Arousal space for CLIP and the ViT, separated by semantic category. Highlighted regions indicate significant clusters where the error increased under cross-category extrapolation (permutation test, cluster-corrected alpha = 0.05). While the Object category shows a shared vulnerable region for both encoders, the peak error locations in the Animal, Person, and Scene categories diverge substantially, indicating model-specific representational gaps.

### 3. Error Vector Field and Occupancy Dynamics Analysis

#### *3.1. Visual Characterization of Error Topology*

To characterize the geometric structure of prediction failures, we constructed error vector fields and occupancy maps for each semantic category. As illustrated in Figure S4, the topological differences between the two encoders are stark. The ViT baseline exhibits a "sink-centric" topology where error vectors point sharply toward a limited number of localized regions. This behavior creates high-occupancy clusters that effectively trap the model's affective output within narrow subspaces. In contrast, CLIP exhibits a more expansive and complex field. Rather than converging toward single points, its vectors often form rotational or divergent patterns, particularly in the Person and Scene categories. This structure prevents trajectories from collapsing into localized sinks, maintaining the representational diversity of the affective mapping.

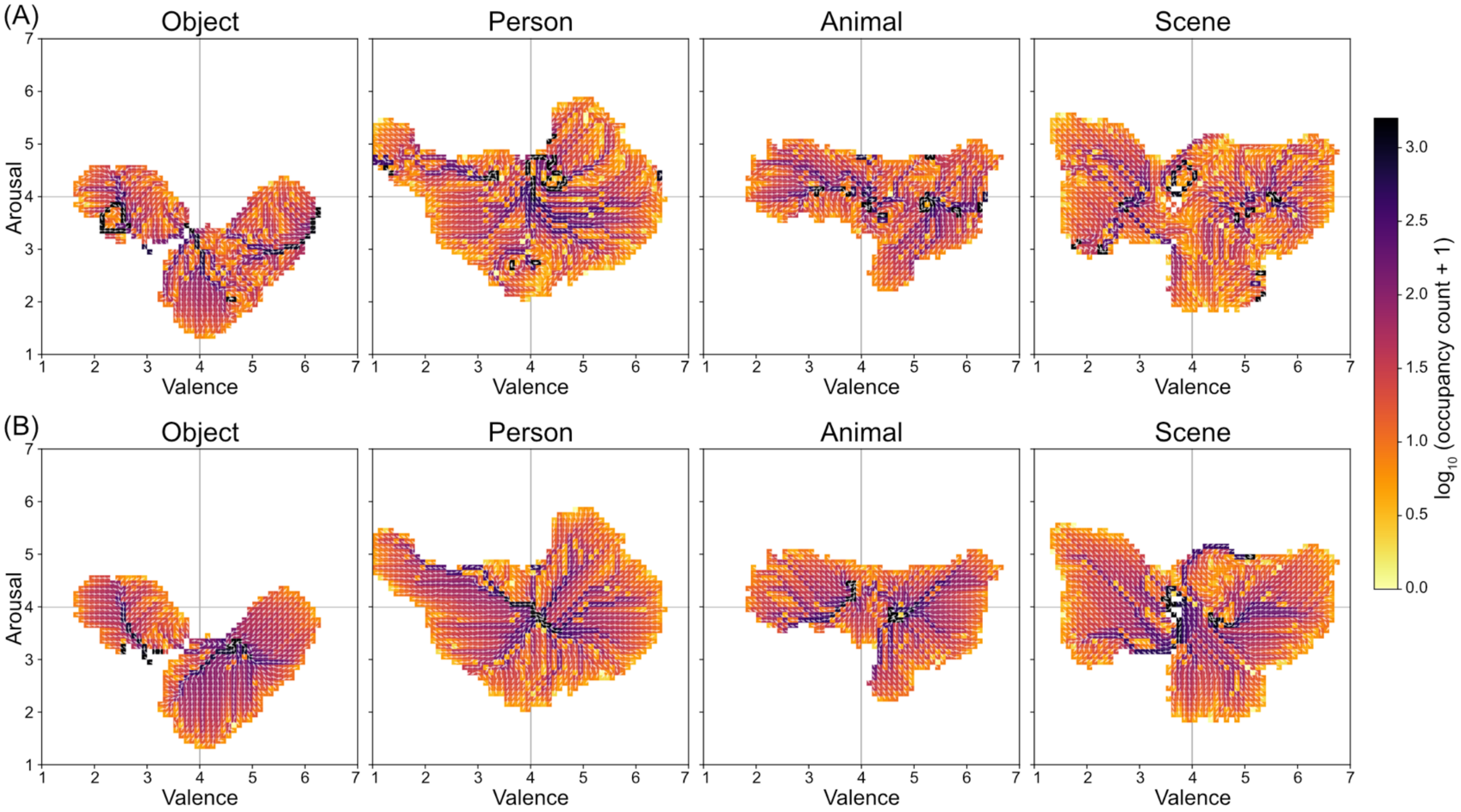


**Figure S4. Category-specific error vector fields and trajectory occupancy maps.** The maps visualize the error dynamics for (A) CLIP and (B) ViT across the four semantic categories. The quiver plots (white arrows) represent the mean error vectors, indicating the direction and magnitude of prediction bias at each location in the Valence–Arousal space. The background heatmaps indicate the log$_{10}$ occupancy count derived from the stochastic particle simulations. CLIP's fields exhibit more complex, rotational structures that distribute occupancy over a wider area. In contrast, the ViT's fields are dominated by strong attractors (bias sinks), where vectors converge toward narrow, high-occupancy regions.

*3.2. Quantitative Analysis of Spatial Dispersion*

The observed topological differences are quantified using normalized spatial entropy ($H_{norm}$, as summarized in Table S2. Across all categories, CLIP consistently achieved higher $H_{norm}$ values than the ViT, confirming that language-aligned representations are less susceptible to concentrated categorical biases.

**Table S2. Spatial dispersion and dynamic diversity metrics for CLIP and ViT across thematic and categorical folds.**

| | LOTO | | LOCO | |
|---|---|---|---|---|
| | CLIP | ViT | CLIP | ViT |
| All | 0.61 | 0.55 | - | - |
| Object | 0.54 | 0.51 | 0.49 | 0.48 |
| Person | 0.69 | 0.47 | 0.43 | 0.42 |
| Animal | 0.58 | 0.48 | 0.45 | 0.52 |
| Scene | 0.51 | 0.48 | 0.55 | 0.41 |

Note: For the LOCO protocol, prediction errors for each category were obtained from the fold in which that category was held out. The LOCO All row is computed by aggregating image-level errors across all four held-out-category folds. Higher $H_{norm}$ indicates more uniform spatial distribution of prediction errors across the Valence–Arousal plane.

*3.3. Robustness and Parameter Sensitivity of Simulation Results*

To verify that the reported $H_{norm}$ differences were not artifacts of specific simulation parameters, we conducted a multi-parameter sensitivity analysis. We systematically varied the trajectory depth (*max_steps* from 20 to 100), the termination criteria (*stall* threshold from 5 to 20 steps), and the sampling density (*n_particles* from 2,000 to 10,000).

As shown in Figure S5, $H_{norm}$ decreased for both encoders as *max_steps* increased, reflecting the convergence of trajectories toward stable regions of high error concentration. The rank order of the two encoders overlapped at very short trajectory lengths, where particles had not yet escaped their initial random positions, but stabilized as the simulation approached convergence. At the trajectory budget used in the main analysis (*max_steps* = 100), the CLIP advantage in H_norm was reproducible across simulation runs. When trajectory depth was held constant, $H_{norm}$ was insensitive to variations in sampling density and stall thresholds, with $H_{norm}$ shifting by less than 5% across the tested ranges. These analyses indicate that the characterization of the error vector fields was not driven by the specific choice of simulation parameters.

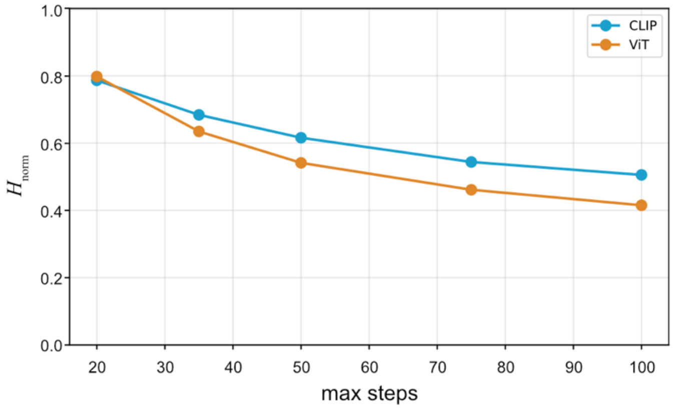


**Figure S5. Sensitivity of spatial dispersion metrics to simulation length.** The line plot illustrates the maximum attainable entropy as a function of the maximum steps per trajectory for CLIP (blue) and the ViT baseline (orange). Data points represent the mean $H_{\mathrm{norm}}$ values derived from 5,000 particles with a stall threshold of 10 steps.

*3.4. The Person Category and Rotational Structure of Error Dynamics*

The largest encoder-level difference in full-model $H_{\mathrm{norm}}$ under LOTO occurred in the Person category, where CLIP exceeded the classification-pretrained ViT by a substantially wider margin than in the other three categories (Table S2). Qualitative inspection of the error vector fields suggests that this divergence reflects not only spatial spread but also a rotational structure in CLIP's error dynamics for Person images. In dynamical systems terms, rotational components in a vector field counteract point-attractors by distributing occupancy mass across a broader manifold or cyclic path, thereby elevating $H_{\mathrm{norm}}$ beyond the levels accounted for by spatial dispersion alone. This suggests that language-aligned representations preserve the multidimensional semantic structure of social affect instead of collapsing it into a singular attractor. The classification-pretrained ViT, by contrast, showed comparatively low $H_{\mathrm{norm}}$ in the Person category, consistent with a representation where predictions are concentrated within narrow regions of the affective space. Notably, this CLIP advantage did not persist under LOCO; when the Person category was withheld, CLIP's $H_{\mathrm{norm}}$ diminished to 0.43. This indicates that while language alignment supports richly structured representations of diverse content, this advantage depends on the presence of category-specific anchors during training. Future work quantifying vector field curl or vorticity would further clarify these qualitative observations.

### 4. Center-oriented Bias by Semantic Category

*4.1. Angular Alignment with the Affective Center*

To determine whether prediction errors were systematically biased toward the center of the affective space (Valence = 4.0, Arousal = 4.0), we analyzed the angular alignment between error vectors and the center-pointing direction. For each image, we computed the cosine similarity and the phase difference between the error vector $\mathbf{e}_i = (\hat{V}_i - V_i,\ \hat{A}_i - A_i)$ and a reference vector from the true rating y to the center (4, 4), i.e., the direction from the true point toward (4, 4). Values near 1 indicate that the prediction error points strongly toward the center along that reference direction. The distribution of these angular differences reveals the degree to which an encoder's failures are "center-oriented." As illustrated in the polar histograms of Figure S6, the ViT baseline displays highly peaked distributions centered at zero phase difference. This indicates that its errors are rigidly and systematically aligned with the direction toward the affective center across all categories. While CLIP also exhibits a center-oriented tendency, its angular distributions are significantly broader and more multi-modal, particularly in the Object and Scene categories. This greater angular spread indicates that CLIP's prediction errors are less constrained by a singular pull toward the neutral center, reflecting a more flexible and semantically grounded internal representation that resists the rigid "collapse" seen in the ViT.

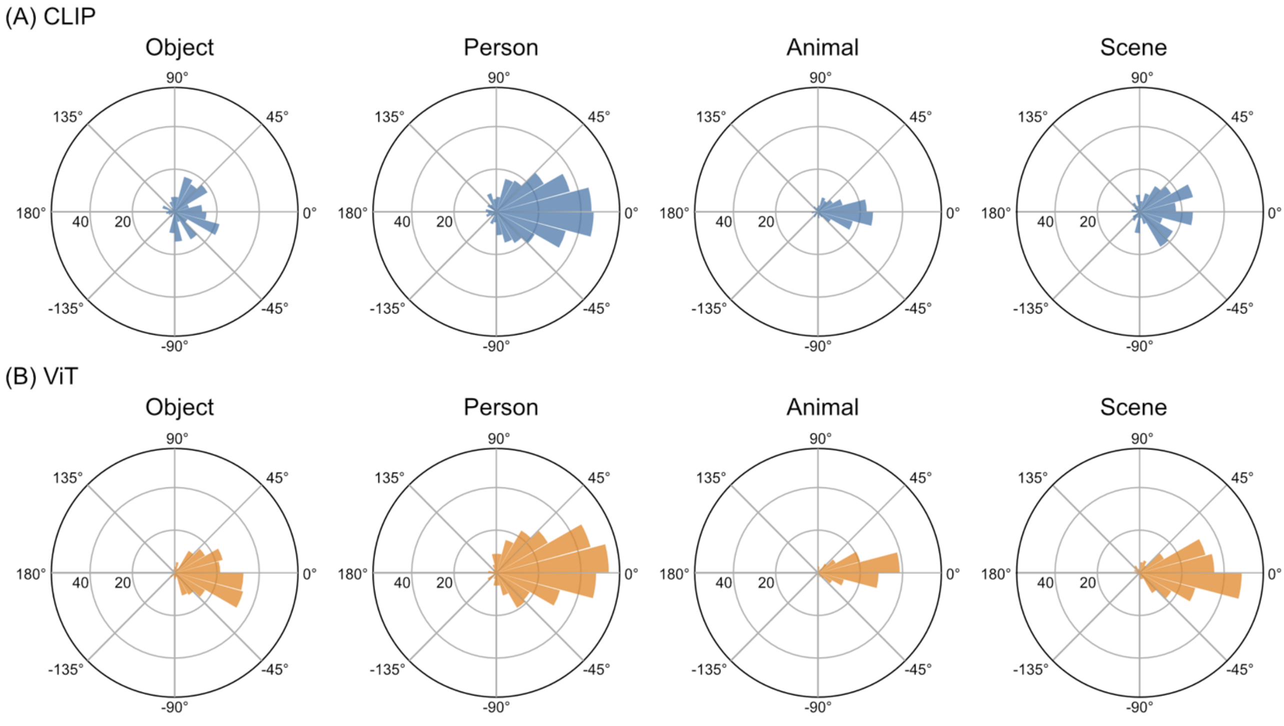


**Figure S6. Angular distribution of error vectors across semantic categories.** Polar histograms illustrate the frequency of error vector directions for (A) CLIP and (B) ViT. Each bin represents the angle of the vector pointing from the predicted state to the true state. The ViT shows narrow, highamplitude peaks, indicating that its prediction errors are highly directional and systematic within each category. CLIP exhibits more distributed angular profiles, particularly in the Person and Scene categories, reflecting a greater diversity in its representational failure modes and a relative lack of rigid categorical bias.

*4.2. Quantitative Summary of Cosine Alignment*

The visual patterns in the polar histograms are quantified using median cosine similarity and effect size measures, as summarized in Table S3. A cosine value near 1.0 indicates that the prediction error points directly toward (4, 4) along the reference direction.

**Table S3. Summary of center-oriented error bias by semantic category**

| | CLIP | ViT | $r_{rb}(\Delta)$ | $d_z$ |
|---|---|---|---|---|
| All | 0.79 | 0.91 | 0.36 | 0.32 |
| Object | 0.62 | 0.89 | 0.60 | 0.53 |
| Person | 0.84 | 0.87 | 0.11 | 0.13 |
| Animal | 0.92 | 0.97 | 0.39 | 0.35 |
| Scene | 0.78 | 0.93 | 0.51 | 0.45 |

Note: $r_{rb}(\Delta)$ and $d_z$ denote the rank-biserial correlation and Cohen's effect size for the paired difference ($\cos_{ViT}$ - $\cos_{CLIP}$), respectively. One-sided Wilcoxon signed-rank tests confirmed that this bias was significant for both encoders across all categories (all $p < 10^{-14}$).

*4.3. Category-specific Patterns and Representational Flexibility*

The strength of this center-oriented bias and the resulting gap between encoders varied markedly across semantic domains. The bias was most pronounced in Object images, which also showed the largest shift toward stronger alignment in the ViT relative to CLIP (*median* $\Delta = 0.17$). In contrast, the Person category showed the smallest gap. While both models attained high median cosines in this domain, the paired difference was only marginally significant ($p = 0.045$). Animal and Scene categories fell between these extremes, with the ViT remaining significantly more center-aligned than CLIP.

In conclusion, every category displayed a significant tendency for errors to point toward the affective center, yet the ViT's alignment was consistently more extreme. CLIP's relative advantage, namely a weaker center-oriented pull, was most evident in domains such as Object and Scene. This suggests that language-aligned pretraining facilitates greater representational flexibility, allowing the model to maintain a more nuanced mapping of affect even when errors occur. The similarity between the two encoders in the Person category further suggests that certain semantic contents may impose stronger shared categorical priors.

**5. Detailed Layer-wise Probing Results by Category**

Linear probing at the category level revealed substantial heterogeneity when Valence and Arousal were examined individually (Fig. S7). At the final layer (layer 12), CLIP's Valence $R^2$ remained negative across all four categories, ranging from -0.53 (Object) to -0.04 (Animal), with a cross-category spread of 0.49. The classification-pretrained ViT showed a different profile at the same layer, with $R^2$ ranging from -0.11 (Animal) to +0.12 (Scene), a spread of 0.23. The ViT yielded positive $R^2$ for Valence in the Person and Scene categories, whereas CLIP's $R^2$ remained negative across all categories. Arousal showed greater cross-category variability than Valence for both encoders. At layer 12, CLIP's Arousal $R^2$ ranged from -1.62 (Object) to -0.51 (Scene), a spread of 1.11. The classification-pretrained ViT's Arousal $R^2$ at the same layer ranged from -1.60 (Object) to -0.25 (Scene), a spread of 1.35. Cross-category spread in Arousal was therefore larger than in Valence for both encoders, with the classification-pretrained ViT showing the largest spread overall. Across both Valence and Arousal at layer 12, the classification-pretrained ViT exhibited greater between-category variability in single-layer $R^2$ than CLIP. This pattern was most evident for Valence, where the ViT's $R^2$ shifted from negative to positive across categories, in contrast to CLIP, which showed consistently negative but more uniform $R^2$ across categories.

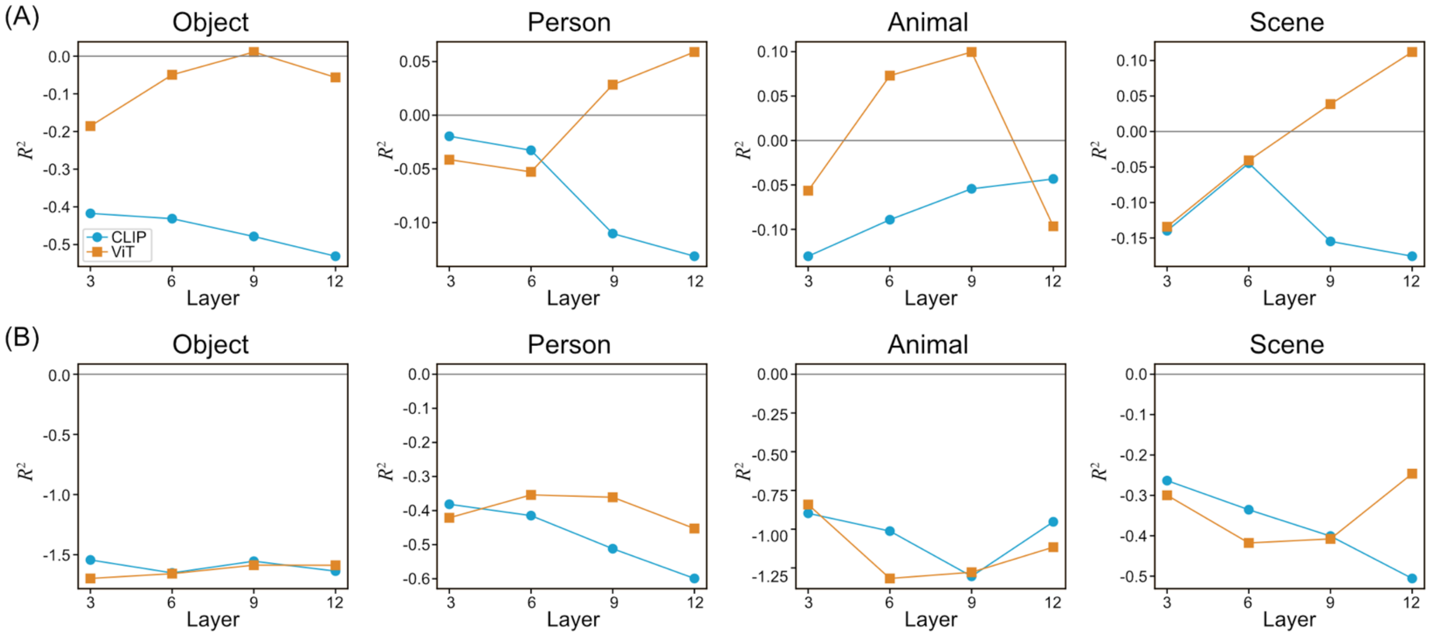

**Figure S7. Layer-wise linear probing performance across semantic categories.**

The plots illustrate the out-of-sample prediction performance $R^2$) for (A) Valence and (B) Arousal as a function of encoder depth at layers 3, 6, 9, and 12. Blue circles represent CLIP, and orange squares represent the ViT baseline. Results are shown for the four semantic categories: Object, Person, Animal, and Scene. The horizontal grey line indicates $R^2$ = 0. While CLIP maintains relatively stable, negative $R^2$ values across most categories, the ViT exhibits significant volatility, including a shift to positive predictive performance for Valence in the Person and Scene categories. Arousal performance shows higher sensitivity to semantic content and overall lower predictive stability for both models compared to Valence.

### 6. Layer-wise $H_{norm}$ Profile and Center-Ward Bias Profiles

#### *6.1. Methods*

$H_{norm}$ and the center-ward cosine alignment were computed at four intermediate Transformer layers (layers 3, 6, 9, and 12) using the same grid-discretization, weighted phase-locking, and trajectory simulation procedures described in Error Vector Field and Dynamics Analysis in the main Methods. For each layer, the layer-$l$ Ridge probe described in Layer-Wise Probing in the main Methods was used to obtain Valence and Arousal predictions under both LOTO and LOCO protocols, and per-image error vectors were aggregated to estimate $H_{norm}$ and the mean cosine similarity between each error vector and the vector pointing toward the geometric center of the affective scale (center-ward bias). As stated in Layer-Wise Probing in the main Methods, the layer-wise values are not directly comparable in absolute scale to the full-model values, since the layer-wise probes use [CLS] token features from intermediate layers and use layer-specific Ridge regularization.

#### *6.2. Pooled Layer-wise $H_{norm}$ and Center-Ward Bias*

At the pooled level (all images aggregated, ignoring category), $H_{norm}$ did not increase or decrease monotonically with depth for either encoder under either protocol (Fig. S8). Under LOTO, CLIP and the classification-pretrained ViT exhibited comparable $H_{norm}$ at L3 (CLIP 0.495 vs. ViT 0.492), with the classification-pretrained ViT exceeding CLIP at L6 (0.505 vs. 0.445) and CLIP exceeding the classification-pretrained ViT at L9 (0.513 vs. 0.427) and L12 (0.564 vs. 0.503). Under LOCO, encoder differences in $H_{norm}$ were uniformly small across all layers ($|\Delta| \leq$ 0.032; CLIP–ViT differences at L3: +0.006; L6: -0.032; L9: +0.003; L12: -0.001), consistent with the protocol-induced reduction in encoder separation observed at the full-model level.

The mean cosine similarity between layer-wise error vectors and the center-ward direction under LOTO was positive at every layer for both encoders, consistent with a regression-to-the-mean tendency in single-layer predictions. The classification-pretrained ViT exceeded CLIP at L3 (0.377 vs. 0.353), with the relationship reversing at L6, L9, and L12, where CLIP exceeded the classification-pretrained ViT (CLIP–ViT differences of approximately +0.06, +0.04, and +0.08, respectively). This pattern differs from the full-model result, in which the classification-pretrained ViT exhibited a stronger center-ward bias than CLIP overall.

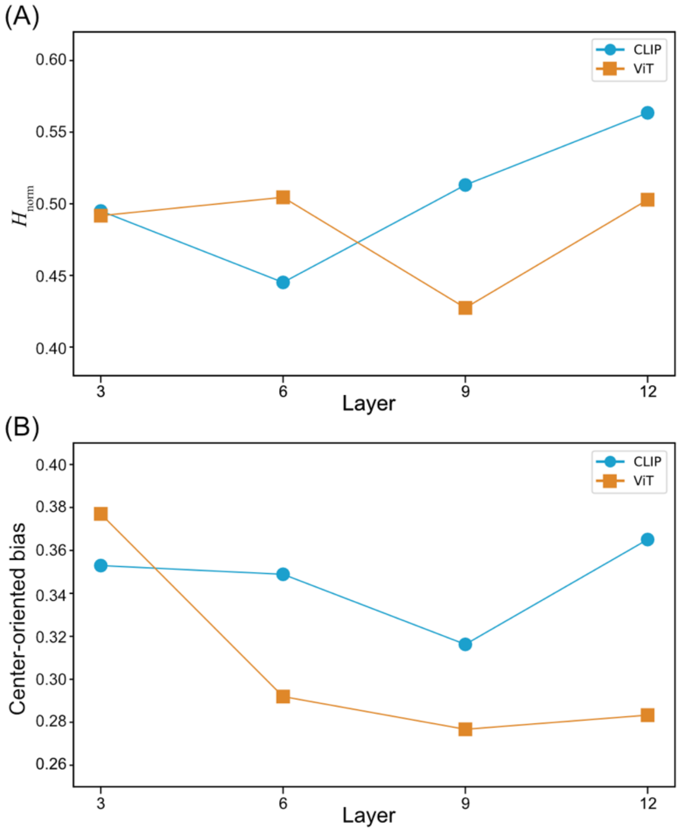


**Figure S8. Pooled layer-wise representational geometry and prediction bias. (A) Normalized spatial entropy across network depth.** The line plots show $H_{norm}$ calculated from error vector fields derived from intermediate layers (3, 6, 9, and 12) pooled across all semantic categories. Solid lines represent the LOTO protocol, while dashed lines represent the LOCO protocol. Neither CLIP (blue) nor the classification-pretrained ViT (orange) exhibits a strictly monotonic trend. **(B) Center-ward bias (mean cosine similarity) across network depth.** The plots show the mean cosine similarity between the layer-wise error vectors and the direction pointing toward the geometric center of the Valence–Arousal space. While both encoders display a consistent positive regression-to-the-mean tendency across all layers, their relative relationship fluctuates.

#### *6.3. Category-Resolved Layer-wise* $H_{norm}$

Category-resolved $H_{norm}$ profiles revealed that the encoder ranking varied across layers within each category, rather than following a category-wide trend (Fig. S9). Under LOTO, no category showed a uniform encoder advantage across all four layers; the most consistent pattern was in the Animal category, where the classification-pretrained ViT exceeded CLIP at three of four layers (L6, L9, L12). The remaining categories showed mixed orderings across depth, with encoder advantage shifting at one or more layers. Under LOCO, encoder differences in $H_{norm}$ were generally smaller than under LOTO, and the layer-wise patterns differed from those observed under LOTO. The most consistent LOCO pattern was in the Person category, where CLIP exceeded the classification-pretrained ViT at

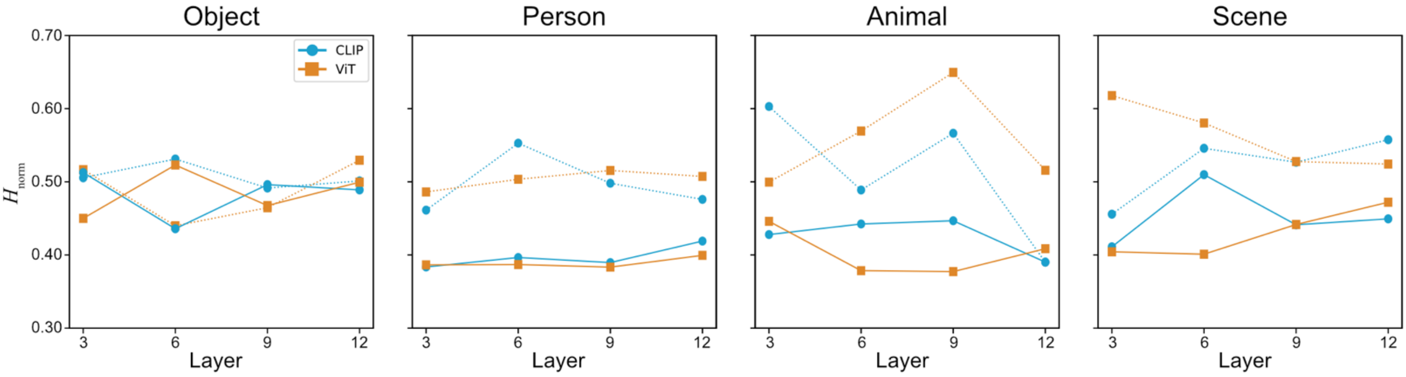


three of four layers (L6, L9, L12). Across categories and protocols, the layer-wise $H_{norm}$ profiles thus show neither a consistent encoder ordering nor a consistent depth trend, indicating that at intermediate layers the relationship between pretraining objective and the spatial dispersion of prediction errors is jointly modulated by category and protocol.

**Figure S9. Category-resolved layer-wise normalized spatial entropy.** The line plots illustrate the trajectory of $H_{norm}$ across intermediate network layers (3, 6, 9, and 12) for CLIP (blue) and the classification-pretrained ViT (orange), separated by semantic category under LOTO (dashed) and LOCO (solid) protocols.

*6.4. Interpretation*

The layer-wise $H_{norm}$ and center-ward bias profiles indicate that the spatial dispersion and directional structure of prediction errors at intermediate Transformer layers depend jointly on the semantic category, the cross-validation protocol, and network depth. The clearer encoder-level differences in error topology emerge at the level of the encoder's final output (full-model analysis, Error Vector Field and Dynamics Analysis in the main Results) rather than at intermediate layers. This pattern is consistent with the view that the language-aligned pretraining objective shapes the geometric structure of affective representations primarily at the encoder's final output, while at intermediate layers the relationship between pretraining objective and error topology is mediated by category-specific factors not detectable in pooled analyses.
